# ReSurveyGermany: Vegetation-plot time-series over the past hundred years in Germany

**DOI:** 10.1101/2022.05.25.493323

**Authors:** Ute Jandt, Helge Bruelheide, Christian Berg, Markus Bernhardt-Römermann, Volker Blüml, Frank Bode, Jürgen Dengler, Martin Diekmann, Hartmut Dierschke, Inken Doerfler, Ute Döring, Stefan Dullinger, Werner Härdtle, Sylvia Haider, Thilo Heinken, Peter Horchler, Florian Jansen, Thomas Kudernatsch, Gisbert Kuhn, Martin Lindner, Silvia Matesanz, Katrin Metze, Stefan Meyer, Frank Müller, Norbert Müller, Tobias Naaf, Cord Peppler-Lisbach, Peter Poschlod, Christiane Roscher, Gert Rosenthal, Sabine B. Rumpf, Wolfgang Schmidt, Joachim Schrautzer, Angelika Schwabe, Peter Schwartze, Thomas Sperle, Nils Stanik, Hans-Georg Stroh, Christian Storm, Winfried Voigt, Andreas von Heßberg, Goddert von Oheimb, Eva-Rosa Wagner, Uwe Wegener, Karsten Wesche, Burghard Wittig, Monika Wulf

## Abstract

Vegetation-plot resurvey data are a main source of information on terrestrial biodiversity change, with records reaching back more than one century. Although more and more data from re-sampled plots have been published, there is not yet a comprehensive open-access dataset available for analysis. Here, we compiled and harmonised vegetation-plot resurvey data from Germany covering almost 100 years. We show the distribution of the plot data in space, time and across habitat types of the European Nature Information System (EUNIS). In addition, we include metadata on geographic location, plot size and vegetation structure. The data allow calculating temporal biodiversity change at the community scale and reach back further into the past than most comparable data yet available. They also enable tracking changes in the incidence and distribution of individual species across Germany. In summary, the data come at a level of detail that holds promise for broadening our understanding of the mechanisms and drivers behind plant diversity change over the last century.

## Background & Summary

The current biodiversity crisis threatens an estimated one million species with extinction ^1^. The nature and rate of observed changes depend on the spatial scale at which they are observed ^2^. At the finest scale, i.e. the local scale of plant communities, vegetation-plot records have been found to become sometimes richer, sometimes poorer in species ^3^, while a considerable temporal species turnover is apparent in the majority of cases ^4^.

Vegetation-plot time series have mainly been collected for particular habitats, such as forests ^5–17^, hedgerows^18^, wet grasslands^19–22^, mesic grasslands^23–29^, dry grasslands^22, 30–35^, acid grasslands and heathlands^36–38^, alpine grasslands^39, 40^, rivers^41^, riverbanks^42^, peatlands^43–46^, roadsides^47^ or arable land^48–50^. Sometimes, they were recorded to assess the changes in species composition across all communities that occur in a certain area^51–55^. So far, vegetation-plot time series have not been accessible without restrictions. In contrast, open access biodiversity time-series data such as BioTIME^56^, comprise all different types of taxonomic groups, ranging from plants, plankton and terrestrial invertebrates to vertebrates, but include only a few vegetation-plot time series. Thus, our database closes a gap for a particular region, which is Germany.

Vegetation-plot resurvey data have been extensively used to assess biodiversity changes by means of monitoring certain vegetation types in local studies, such as managed grassands ^24^and rivers ^41^. More recently, time series have been collected across regions, exploring the contribution of local biodiversity change ^3^ to that observed at broader spatial scales ^1, 57, 58^. While these analyses often failed to detect changes in species richness ^3, 59, 60^, they were able to relate the observed trends to changes in land use and climate ^61, 62^. Although these studies have compiled databases on vegetation-plot time series, they are currently not openly available. This is also the case for the current initiative of ReSurveyEurope, which collates and mobilizes vegetation-plot data with repeated measurements over time (http://euroveg.org/eva-database-re-survey-europe). Our aim is to provide a comprehensive and taxonomically standardised database of vegetation-plot time series for Germany. We confined the geographical extent to Germany because of a long tradition of German vegetation scientists carrying out temporal observations on permanent plots (e.g.^28^), the large amount of available data, our familiarity with the regional literature, and of recent initiatives to mobilize retrospective biodiversity data for trend analyses (www.idiv.de/smon).

Vegetation-plot time series differ in some fundamental ways from other biodiversity time series. Since the advent of phytosociology in the early 20th century^63, 64^, vegetation surveys were carried out in a standardised way. Plot sizes of vegetation relevés can vary considerably and depend on the vegetation type considered (e.g. forest plots usually have plot sizes between 100 and 1000 m^2^, while non-forest plots mostly range from 4 to 100 m^2 65^). In addition, sampling protocols might vary between studies, but they all include complete lists of species occurring at the plot at the time of sampling. In consequence, vegetation-plot records provide information on both presences and absences of species in a community. As sampling is usually done by professionals, absences of a previously occurring species in a time series strongly indicates local extinction, or vice versa, the presence of a species that had not been recorded previously is a robust indication of colonization. However, even with experts carrying out the survey, it is possible that some species may remain undetected in the record because of their phenology or taxonomic uncertainties ^65^. Yet, such vegetation-plot data are much more reliable than vegetation surveys at larger scales, such as floristic grid mapping, where false absence data are the rule ^66, 67^. In contrast to time series at broader spatial scales, vegetation-plot time series contain information on species co-occurrence at scales relevant for direct biotic interactions among individuals ^68^. An additional advantage of vegetation-plot records is that they report the relative abundance of species, in the case of vegetation records from Germany, typically assessed as cover values ^65, 69^. Thus, vegetation-plot records allow testing key theories of biogeography, such as the abundance–range size relationship ^70^ or the relationship between local abundance and niche breadth ^71, 72^. Most importantly, several vegetation-plot time series precede the onset of any other systematic plant species monitoring programme, such as for example the monitoring of Natura 2000 sites in Europe, which only started in 2001^73^. This is particularly important because severe biodiversity loss may have already happened in the second half of the 20^th^ century, mainly brought about by shifts in the type and intensity of landuse as the consequence of technical progress and societal changes ^74^. Finally, species-abundance data in plots can be linked to functional information on species ^65^, which allows the interpretation of the underlying ecological drivers of the changes observed and the consequences for ecosystem functioning^75^.

Based on the data described here we analysed for the first time the dynamics of losses and gains of plant species^76^. We showed that the difference in cover changes between decreasing and increasing species results in biodiversity change even if species richness at the plot scale remains unchanged. Two mechanisms are responsible for these changes. First, losses at the plot scale were more evenly distributed among losing species than gains among winning species. Second, gains and losses in cover were concentrated in different species, resulting in a higher number of losers than winners at the spatial scale of Germany. The temporal extent of the data allowed us to demonstrate that most species losses occurred already in the 1960s, affecting mostly species from mires and spring fens, grasslands and arable land. Thus, these data already helped to shed light on the most important mechanisms underlying biodiversity change in the second half of the 20^th^ century.

## Data Records

ReSurveyGermany is the most comprehensive compilation of repeated long-term vegetation plot records from Germany to date, including published studies as well as surveys from grey literature and nature conservation assessments. A list of all 92 projects included in the study is provided in Table 1. A project might comprise one or several studies and focus on one or several vegetation types, but typically carried out the surveys at the same times and with the same methodology. In total, the projects comprise 1,794 vascular plant species recorded in 7,738 vegetation plots. The plots were either marked with poles or magnets (permanent) or recovered from exact descriptions (semi-permanent). In addition, there were also studies where plots were not matched in time but a set of plots at a site was compared within another set of plots at the same site in the resurvey (community comparison, Fig. 1). We only considered records with complete lists of vascular plants and information on their relative abundance, which was mostly expressed as percentage cover^77^. A further important criterion for including a study was the existence of vegetation data for at least two points in time, although the number of visits (i.e. vegetation records) per site ranged between two and 54. The time span covered by each project is shown in Fig. 1. All records were made between 1927 and 2020. In total, ReSurvey Germany comprises 23,641 vegetation-plot records and 458,311 species cover records.

**Fig. 1:**
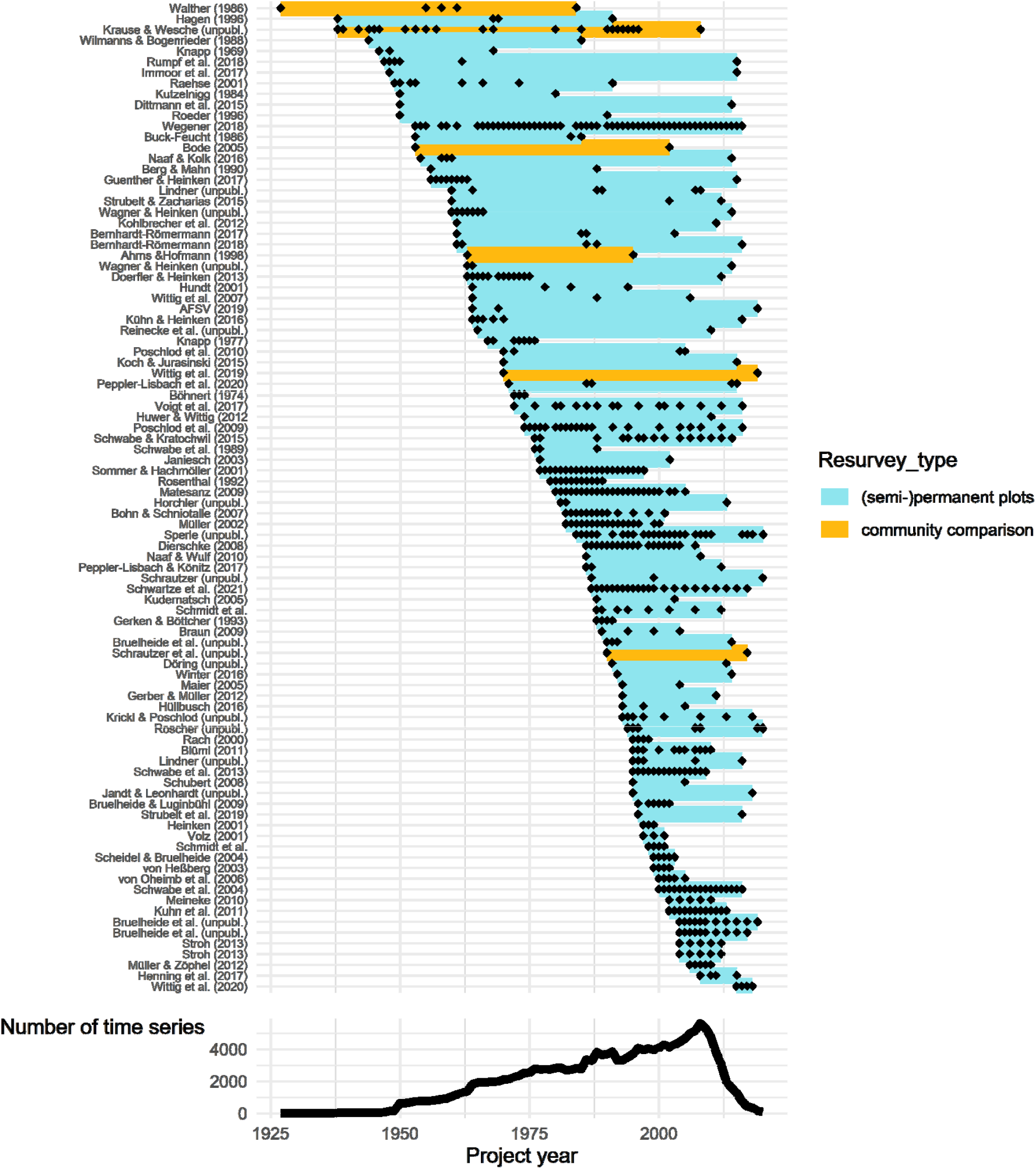
Temporal coverage of the 92 projects included in the study. The coloured lines indicate the start and the end of a project, black diamonds show in which years surveys were made. Resurvey type refers to either studies that were repeated within a particular community across a site without attempts to match plots (community comparison), or were carried out on matched plots, which were either permanently marked or retrieved from exact descriptions (semi-permanent). The lower graph shows the number of times a particular year was included in the covered time span of any of the projects. For a list of projects see Table 1.

**Table 1:**
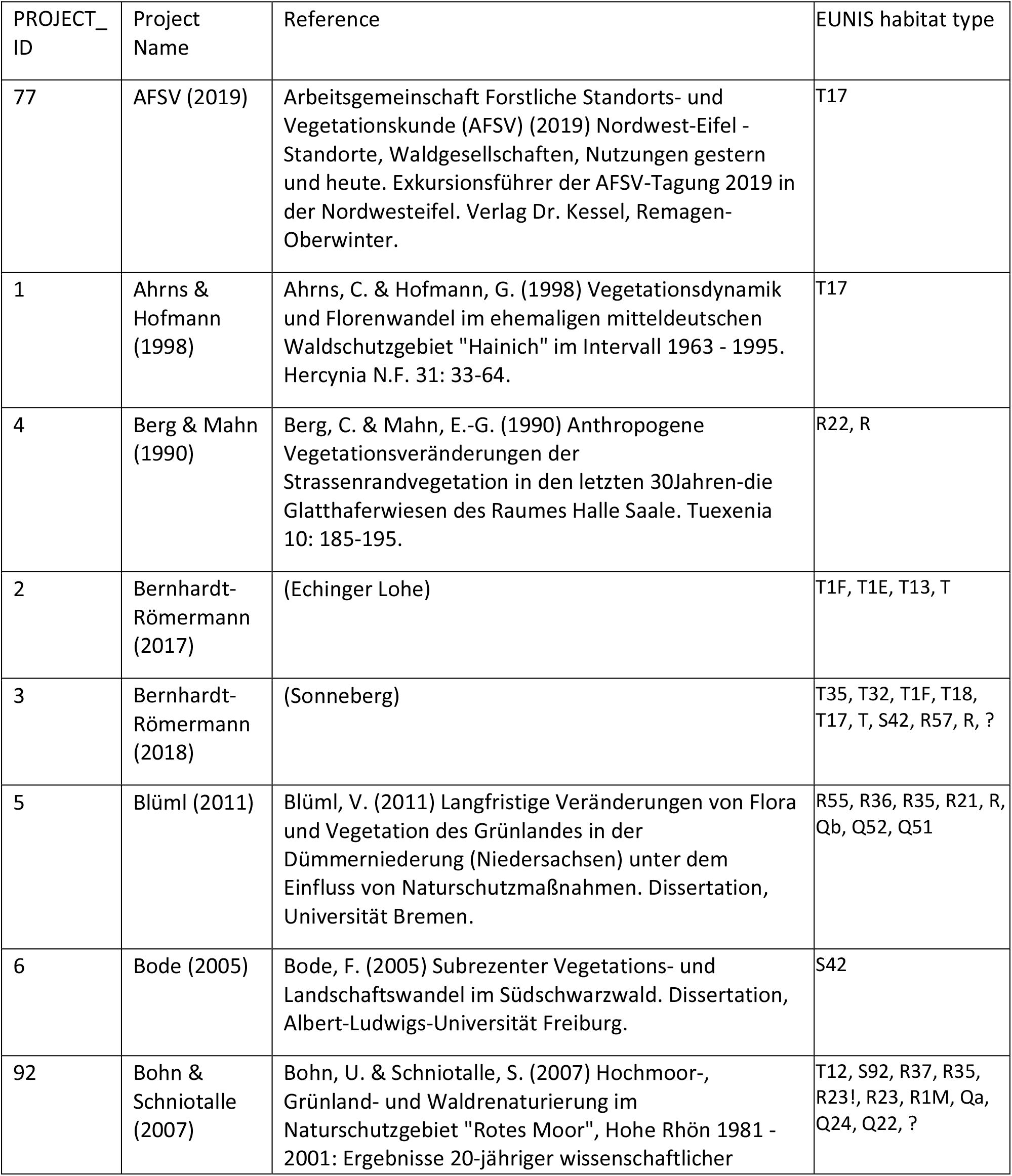
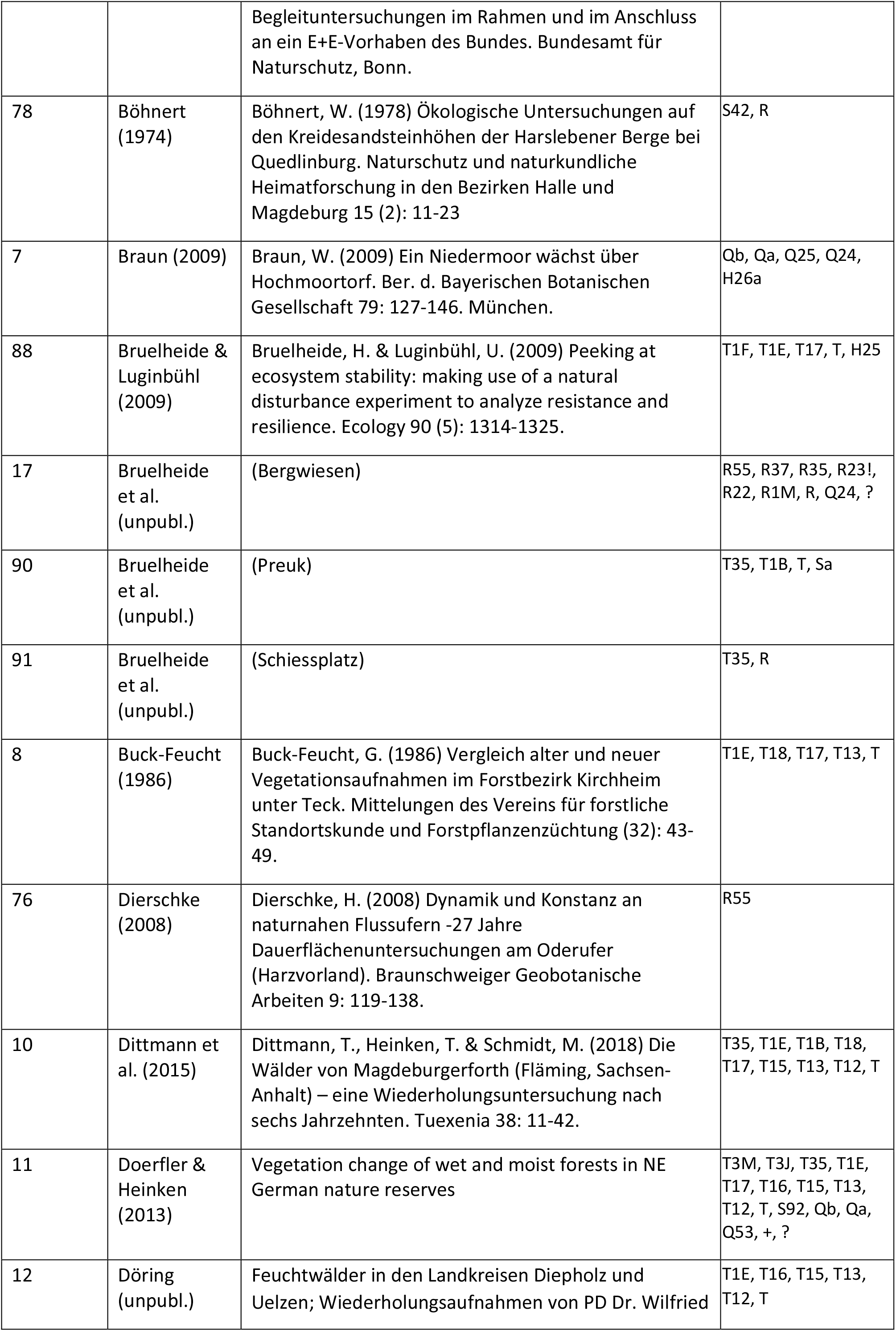
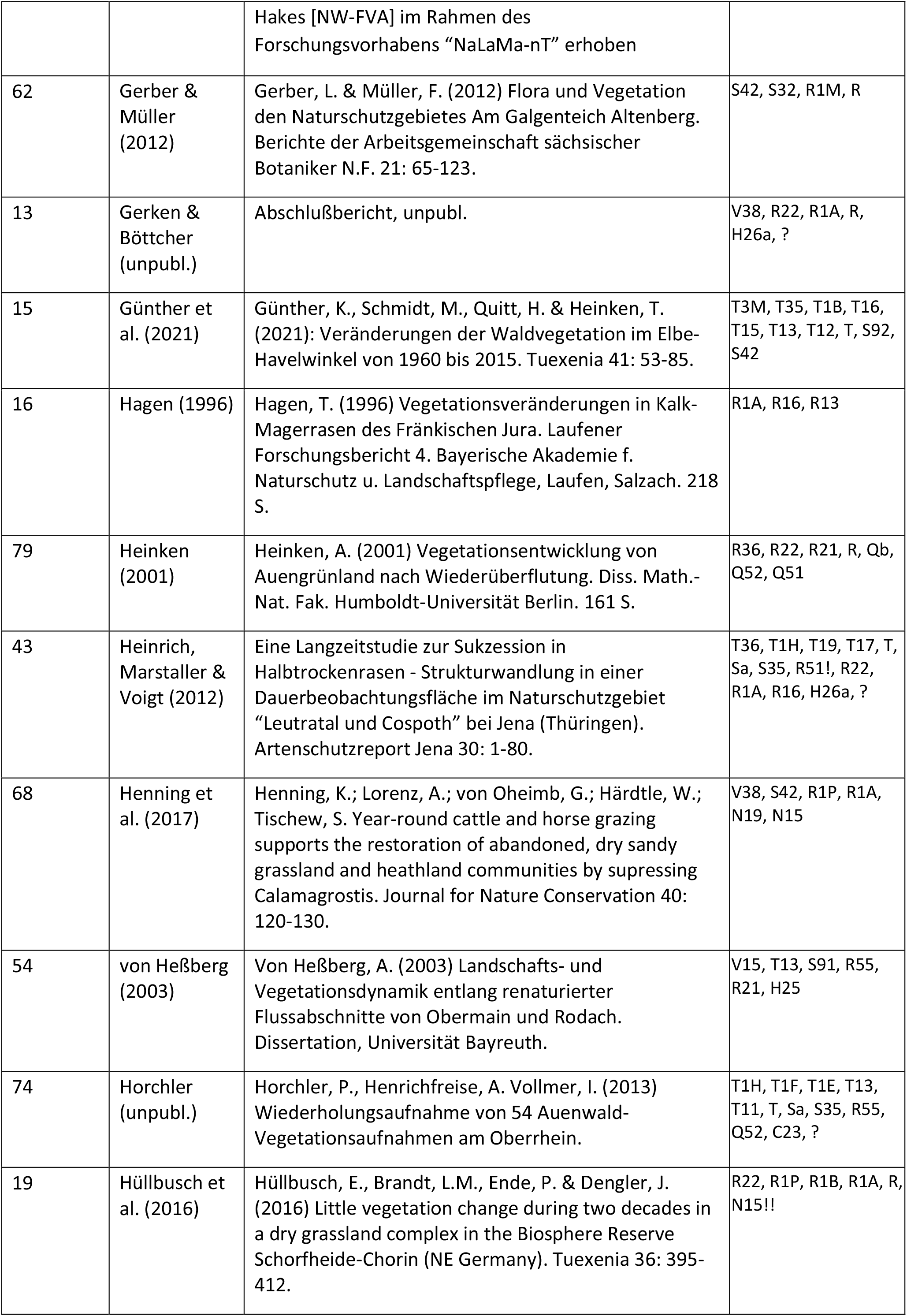
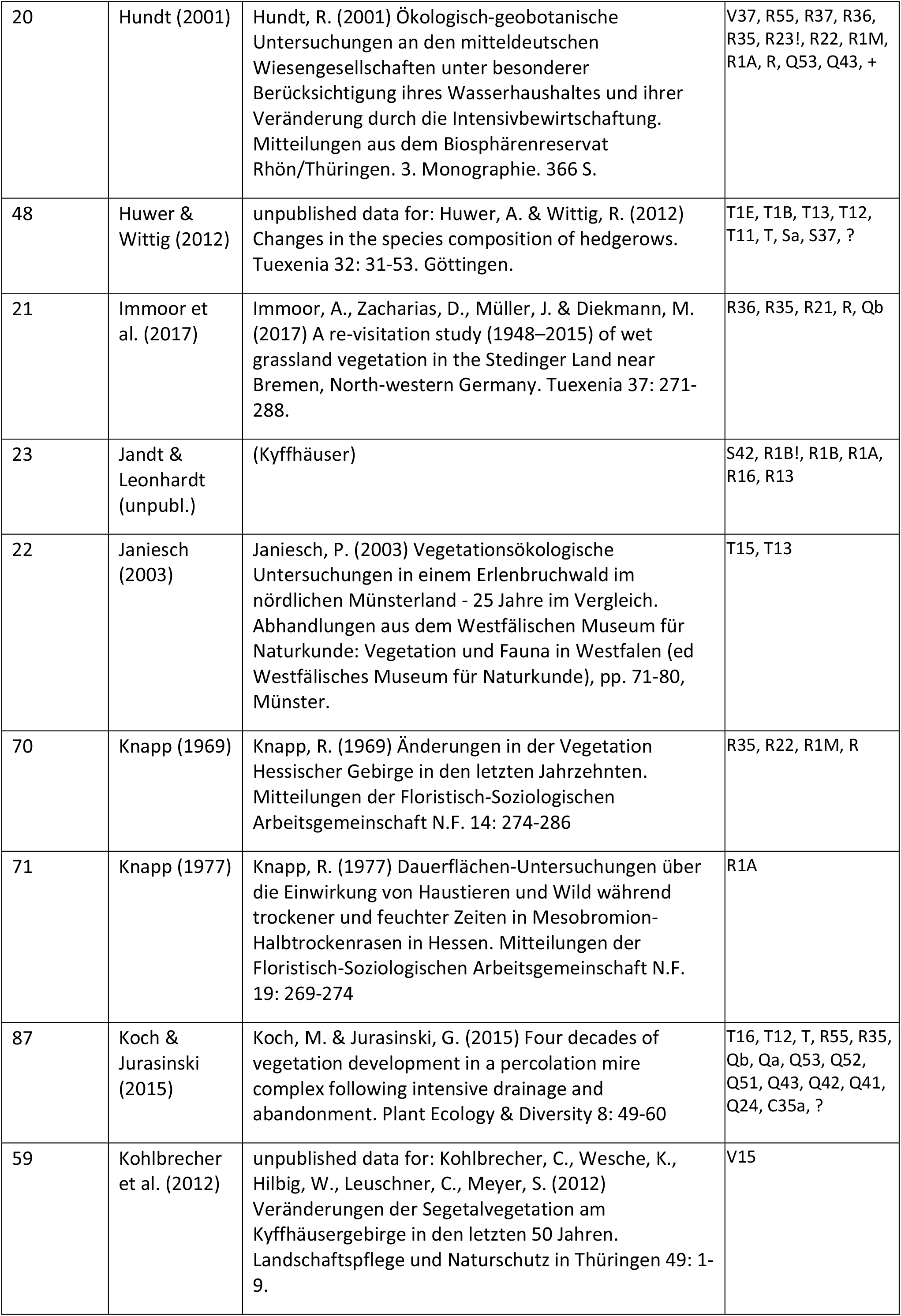
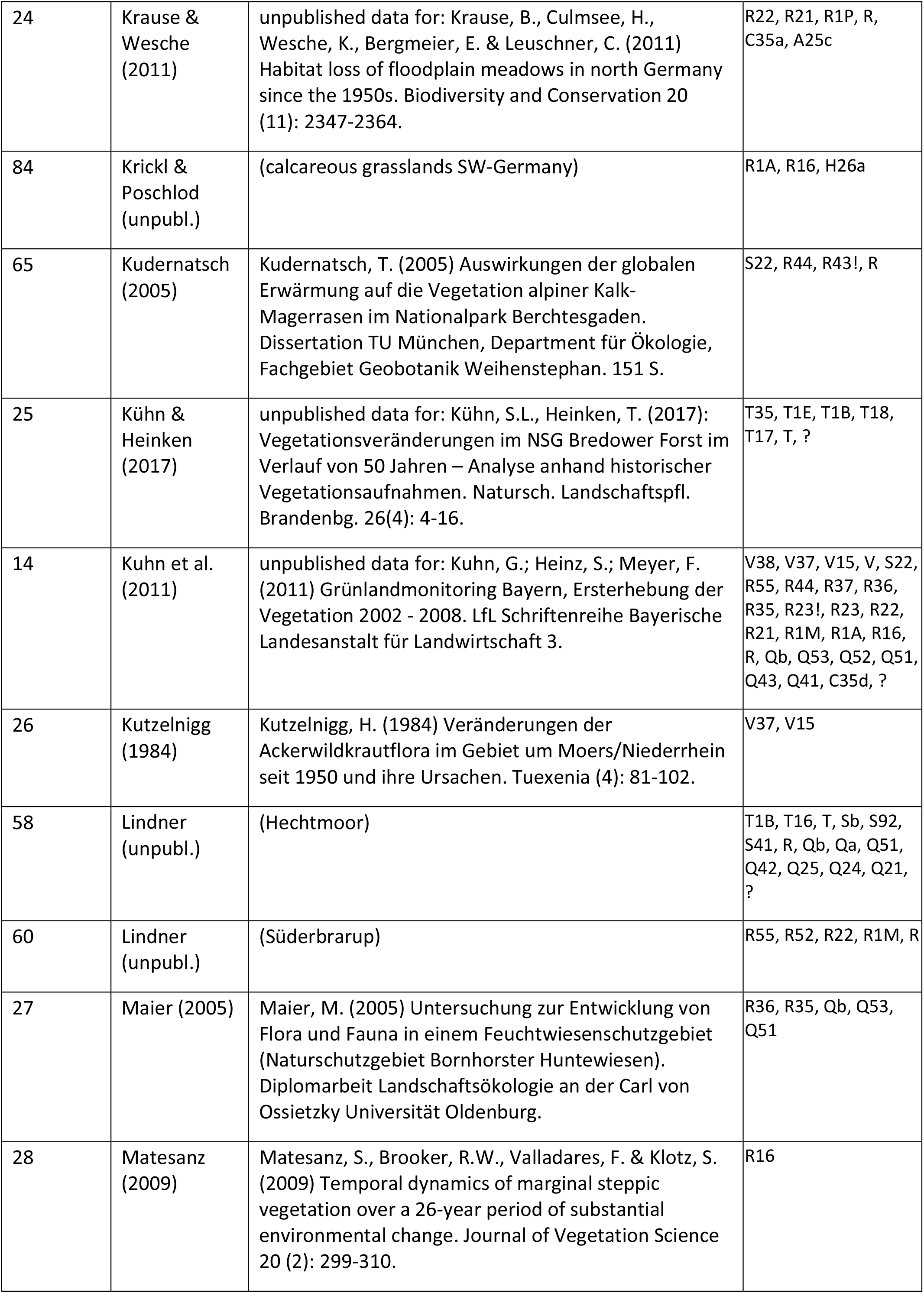
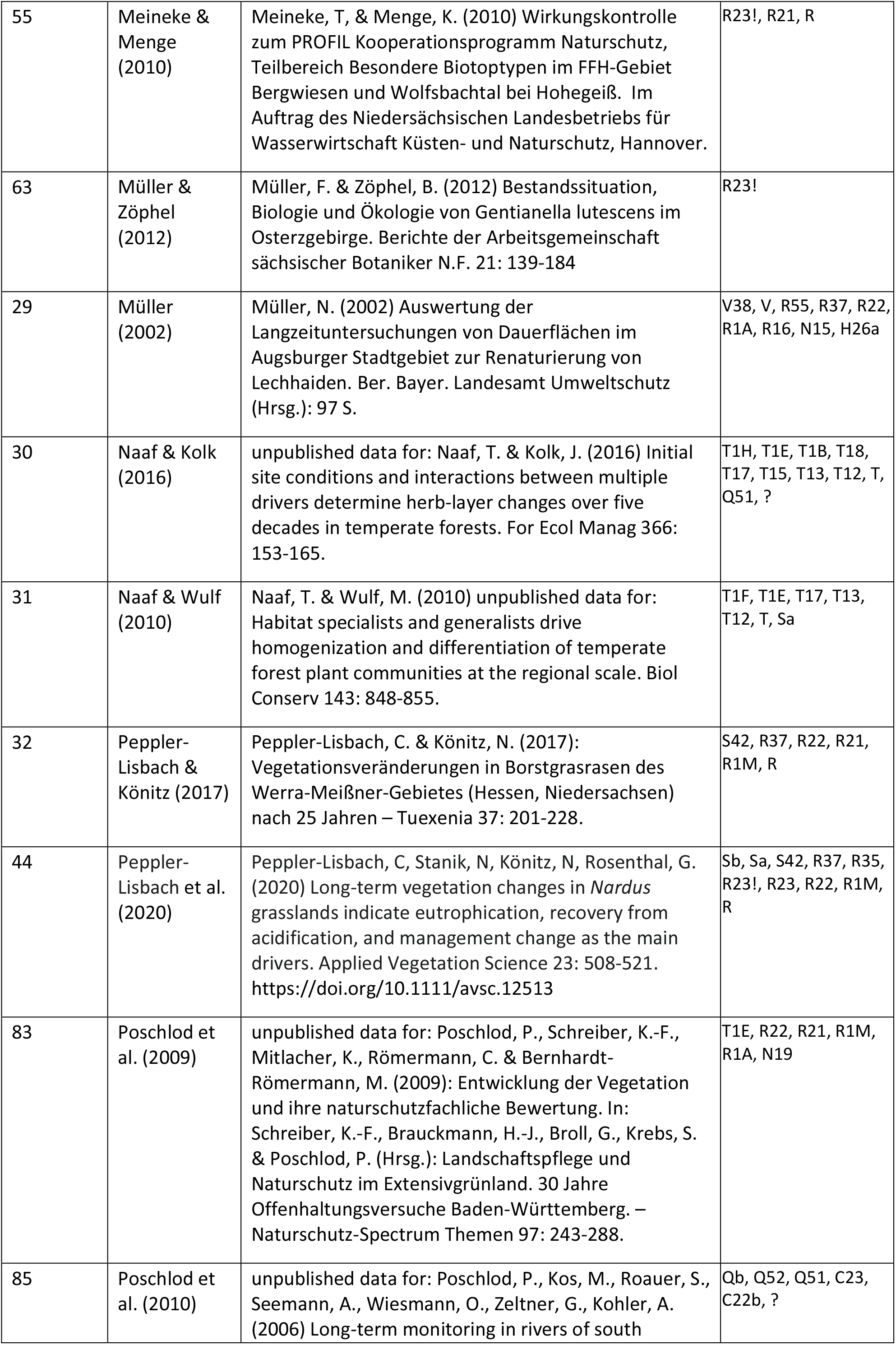
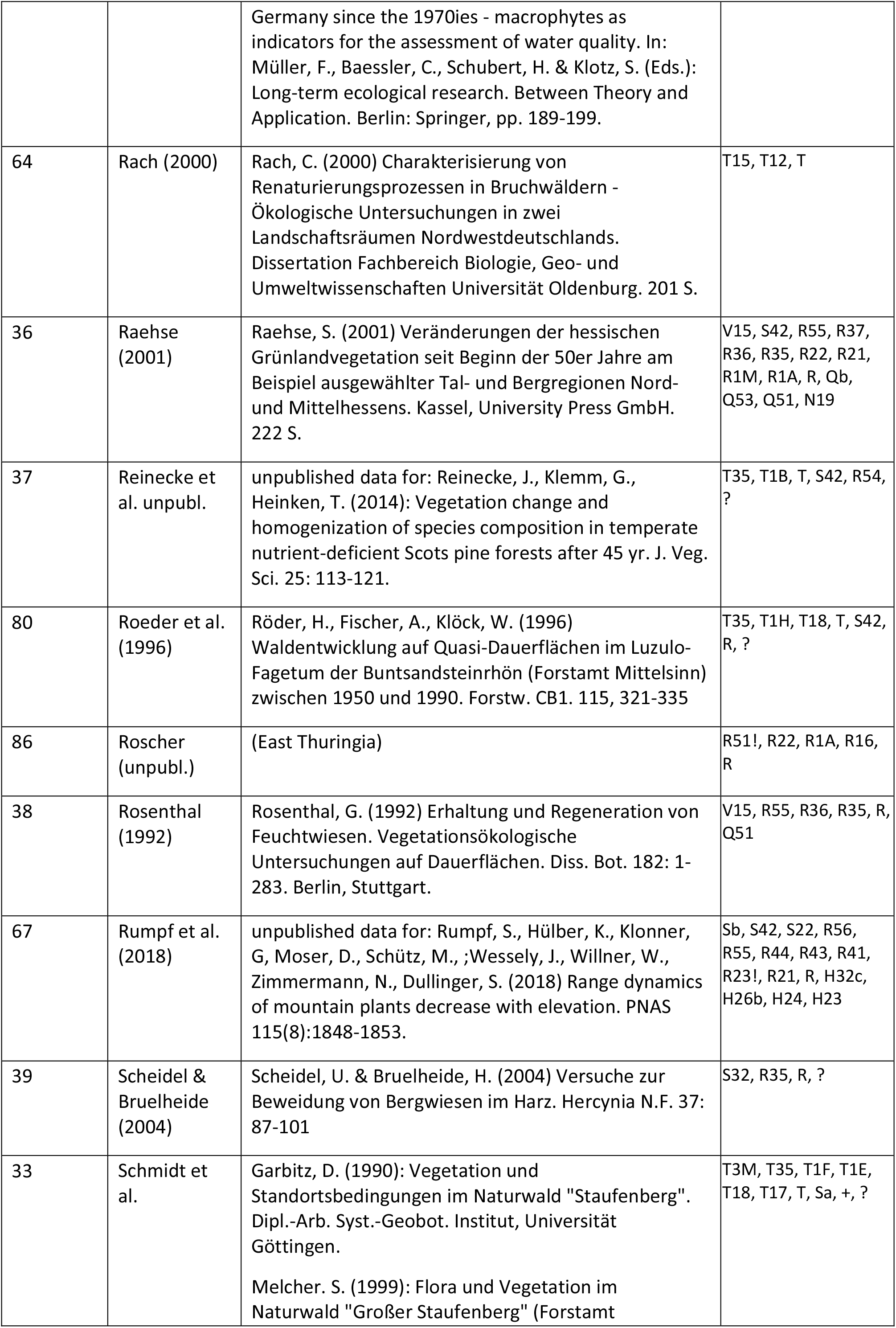
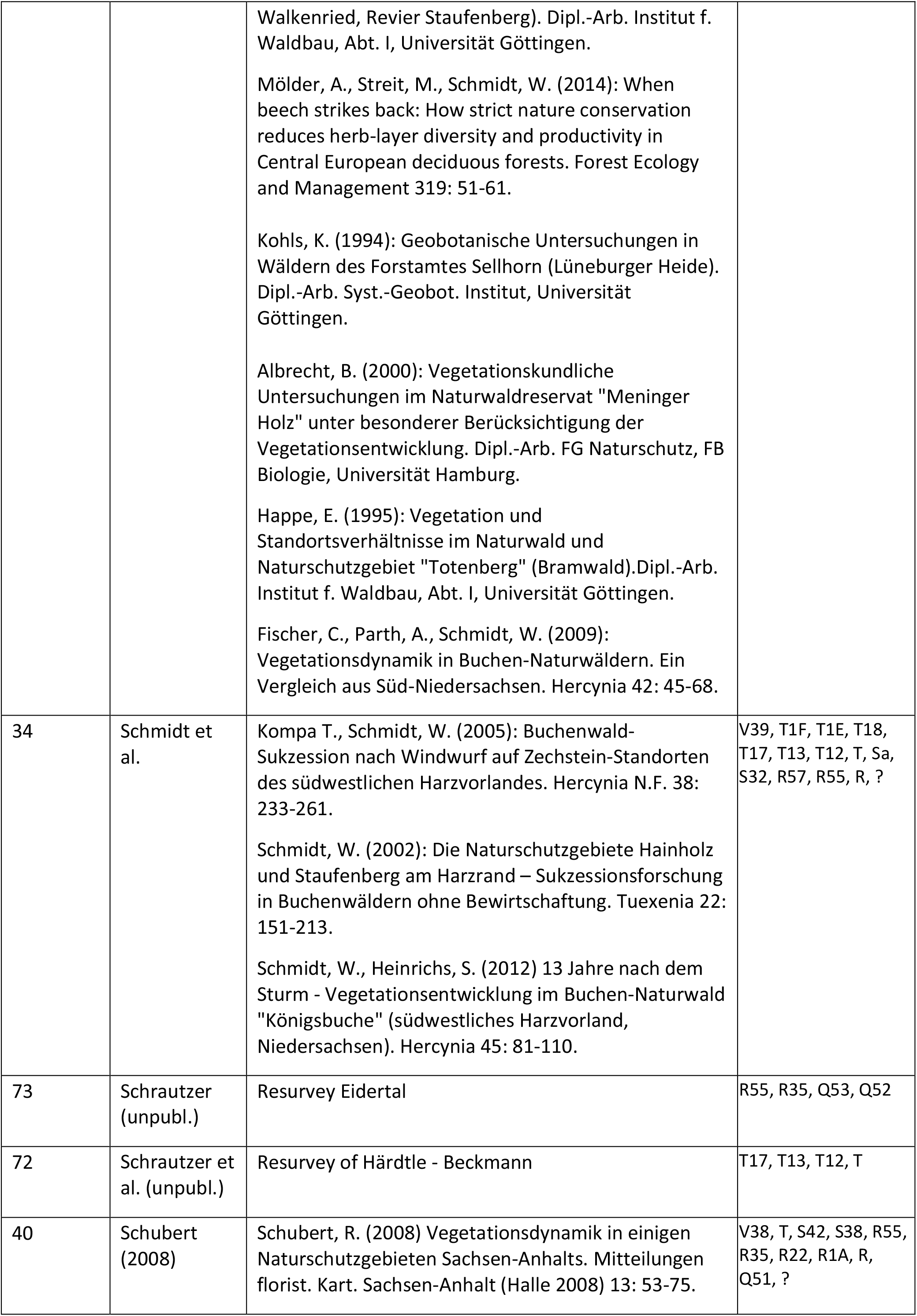
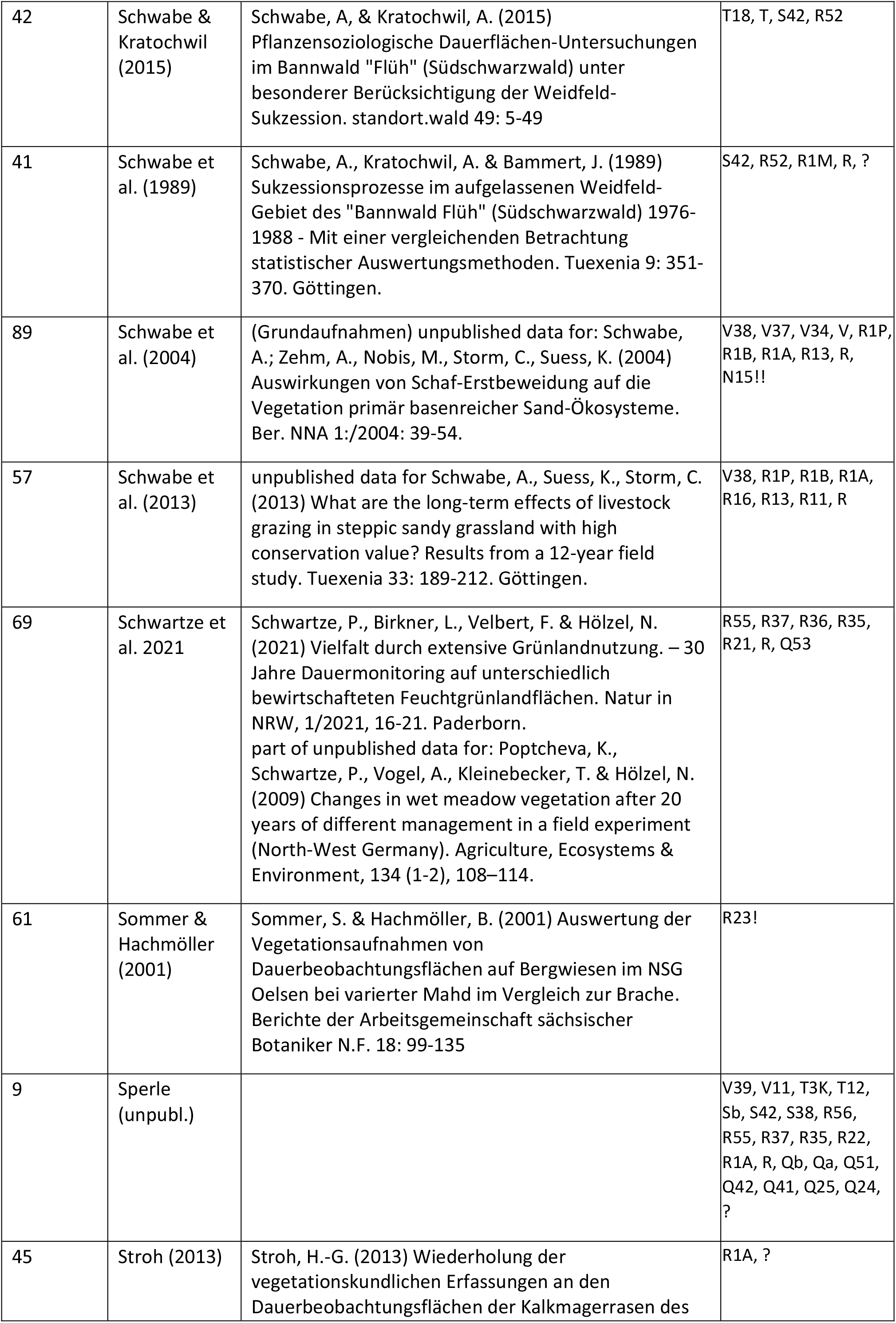
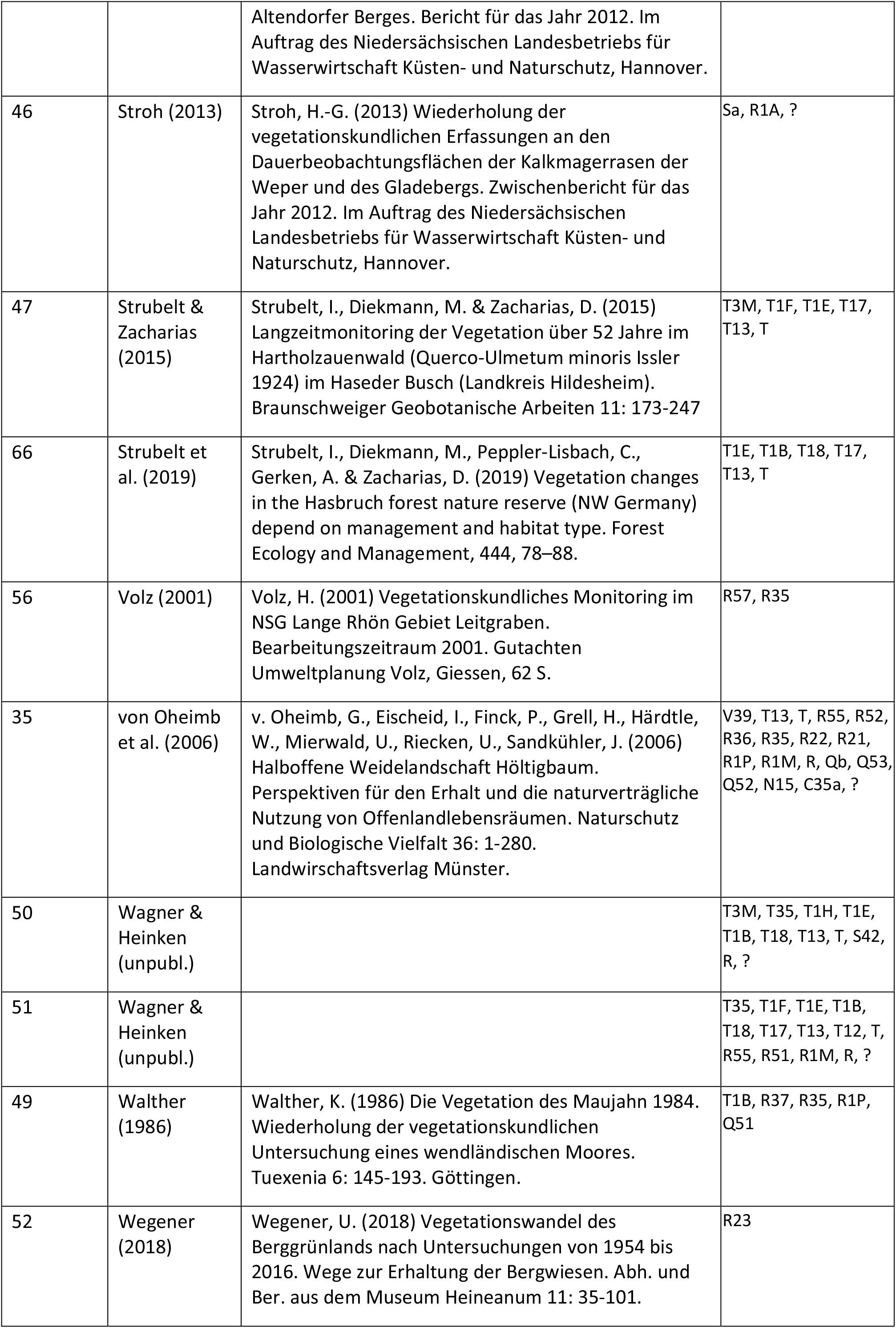
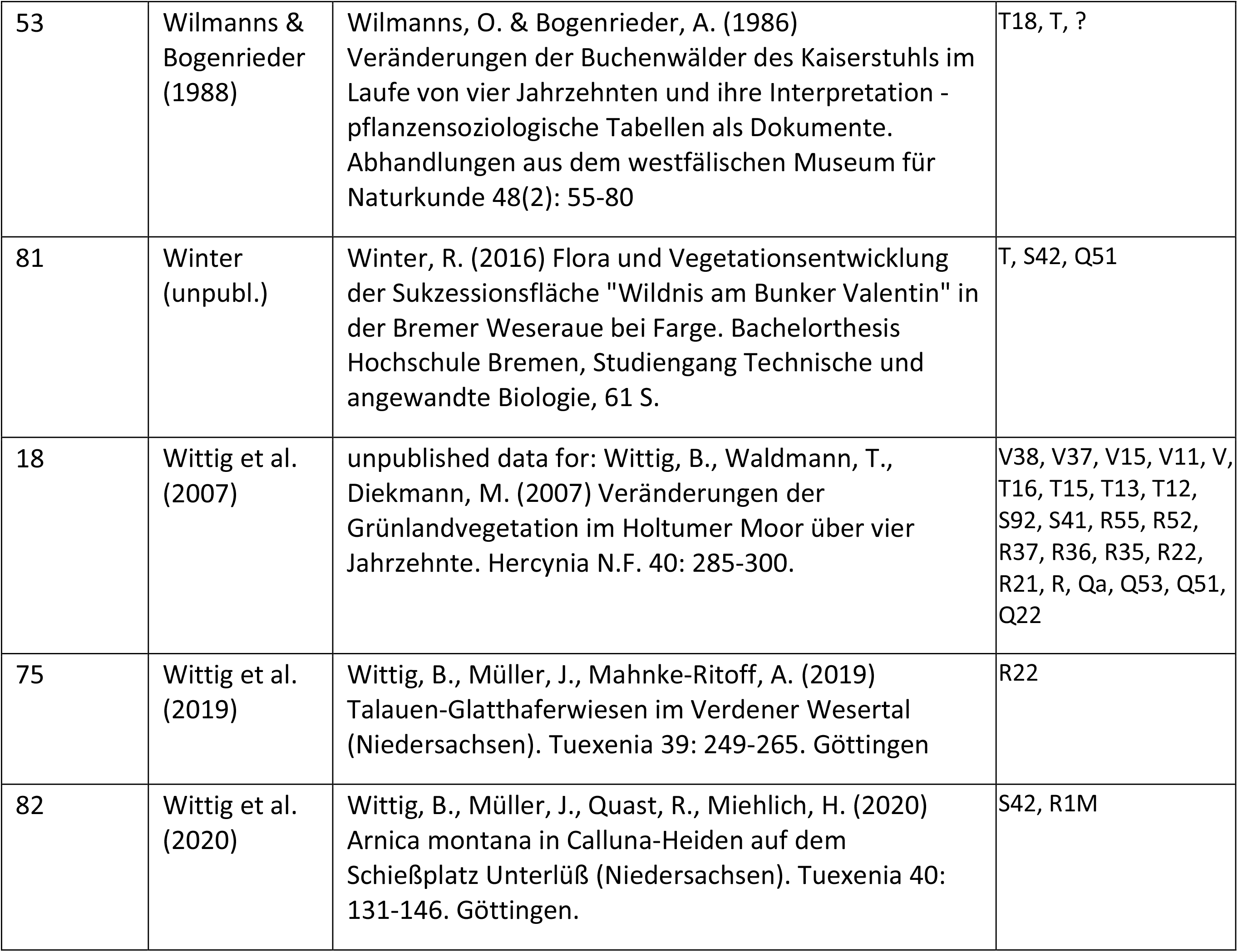
List of all projects included in this study. PROJECT_ID: internal reference number. EUNIS habitat types of time series were assigned to the habitat type by using the earliest plot record that resulted in level 3 EUNIS classification. The classification was based on the EUNIS-ESy expert system^78^ using the R code implementation^79^. If a project included several habitat types, they are shown in decreasing numbers of plot records. Code for habitat types are ?: plots not assigned to any level 3 EUNIS habitat type, +: assigned to more than one level 3 EUNIS habitat type, A: Marine habitats, C: Inland surface waters, H: Inland sparsely vegetated habitats or devoid of vegetation, N: Coastal habitats, Q: Wetlands, R: Grasslands and lands dominated by forbs, mosses or lichens, S: Heathlands, scrub and tundra, T: Forests and other wooded land, V: Vegetated man-made habitats, including arable land.

Plot locations are not evenly spread across Germany (Fig. 2). We assigned the individual plot locations to the grid cells of the quadrants of German ordnance maps (“MTBQ,” 0°10’ × 0°6’, approximately 5.6 km × 5.9 km in the centre of Germany), and tested whether the grid cells with vegetation-plot time-series records differed from those without observations with respect to population density, road density, urban cover, cropland cover and protected areas. This revealed that the sampled grid cells were not representative for the whole area of Germany. Surprisingly, the sampled grid cells showed significantly higher human population densities, road densities and urban cover, while cover of cropland and the amount of protected area was lower (Table 2), which indicates that the majority of time series was made in regions with higher human pressures. The lack of spatial representativeness also becomes obvious when plotting maps of plot locations by the decade of the times when they were visited (Fig. 3).

**Fig. 2:**
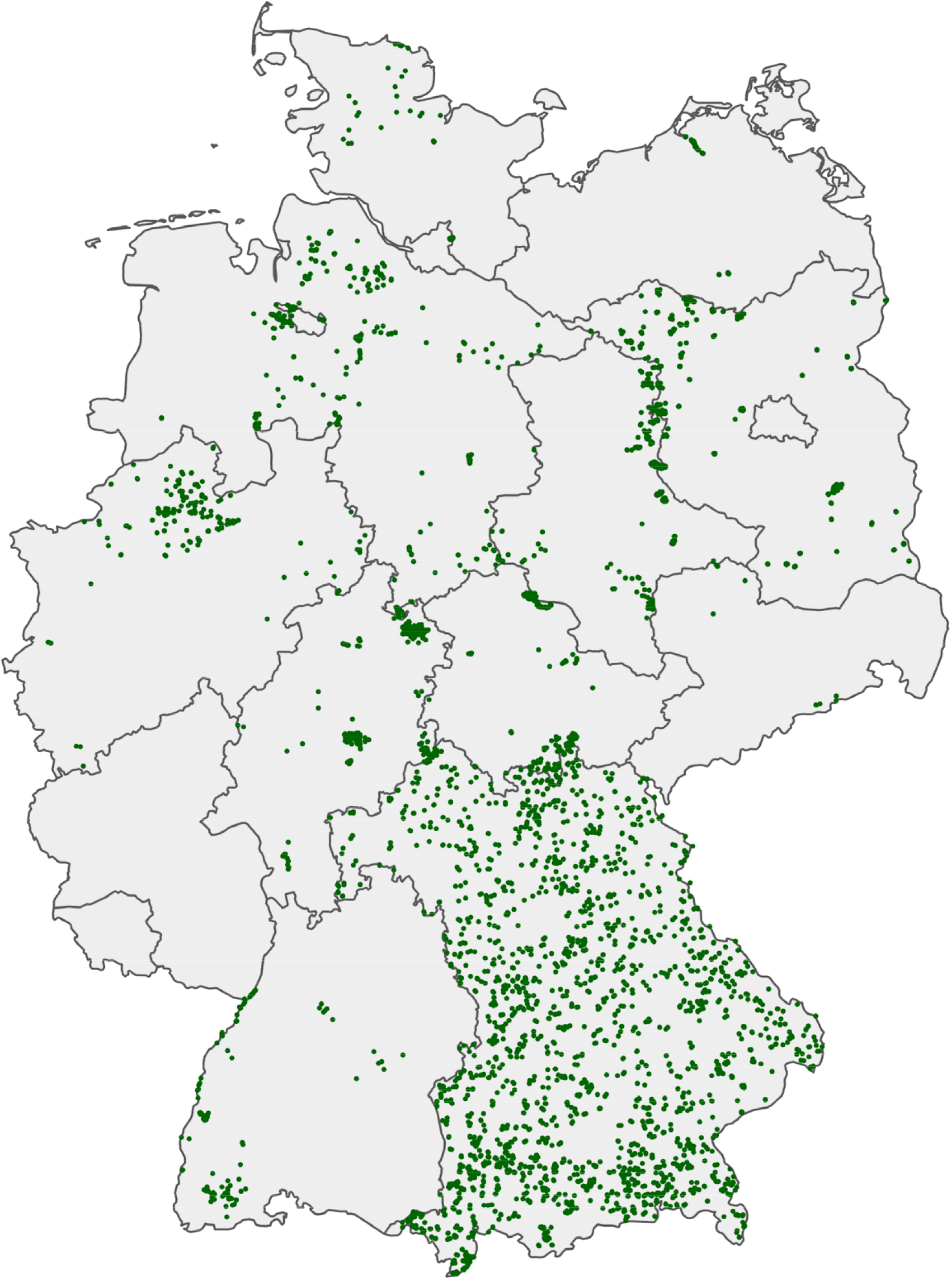
Map of all plots of all projects (n=23,641). Note that green dots may represent one or several plots whih were summarised under the same plot resurvey ID (n=7,738). The more complete coverage of Bavaria resulted from including the grassland monitoring Bavaria which started in 2002^24^.

**Fig. 3:**
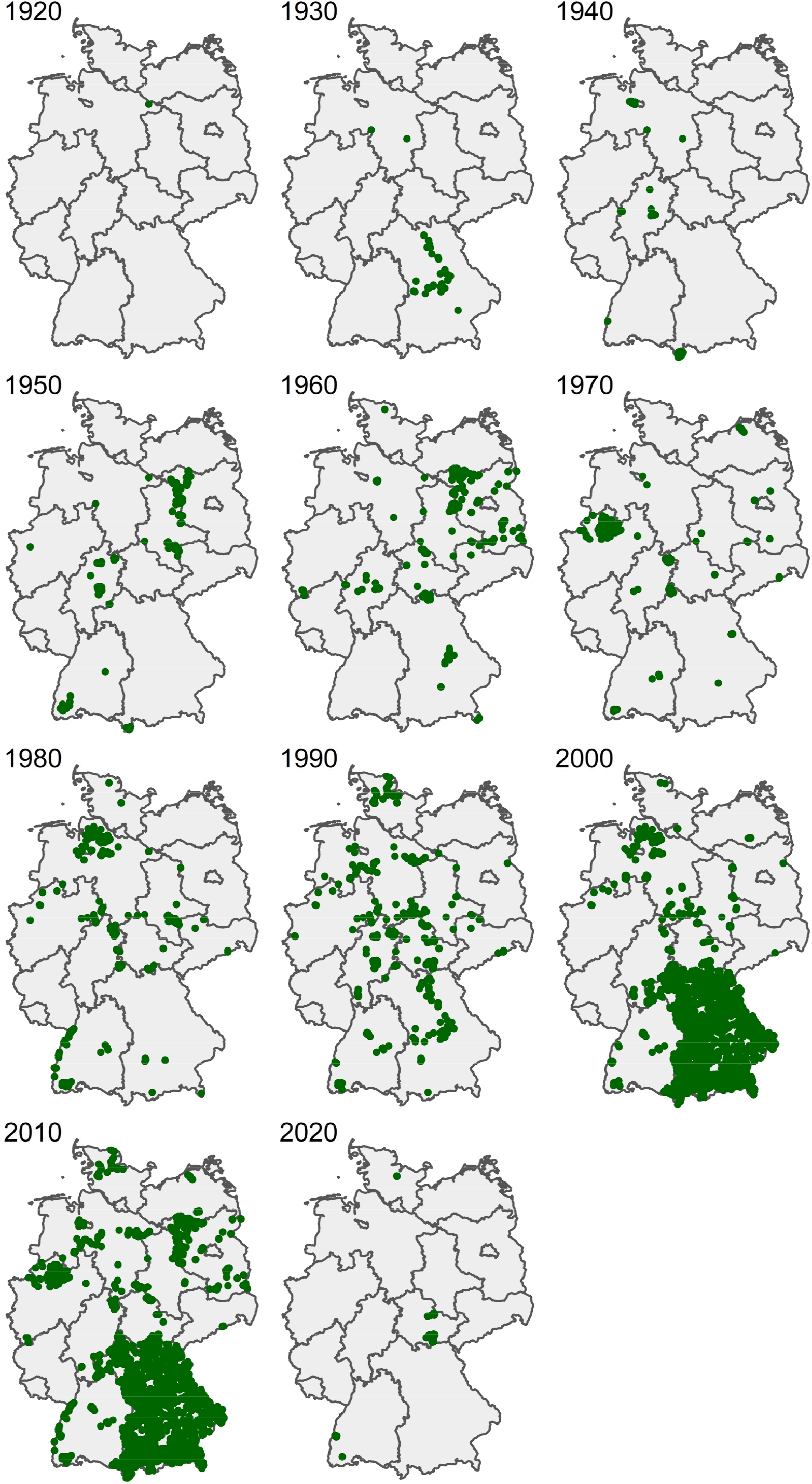
Map of plot visits by decade, with the year showing the begin of the decade.

**Table 2:** Representativeness of grid cells (“MTBQ,” 0°10’ × 0°6’) with time series. The estimates were obtained from linear models comparing samples with unsampled MTBQs with respect to population density, road density, urban cover, cropland cover and protected area.

| Predictors | Population density |  |  | Road density |  |  | Urban cover |  |  | Cropland cover |  |  | Protected area |  |  |
| --- | --- | --- | --- | --- | --- | --- | --- | --- | --- | --- | --- | --- | --- | --- | --- |
|  | Estimates | CI | p | Estimates | CI | p | Estimates | CI | p | Estimates | CI | p | Estimates | CI | p |
| (Intercept) | 30547 | 30128 –<br>309645 | <0.001 | 1.59 | 1.55 –<br>1.63 | <0.001 | 0.07 | 0.06 –<br>0.07 | <0.001 | 0.45 | 0.44 –<br>0.45 | <0.001 | 0.01 | 0.01 –<br>0.01 | <0.001 |
| type<br>[unsampled] | -6686 | -7447 –<br>-5925 | <0.001 | -0.52 | -0.59 –<br>-0.45 | <0.001 | -0.03 | -0.03 –<br>-0.03 | <0.001 | 0.01 | 0.00 –<br>0.02 | 0.029 | 0.01 | 0.01 –<br>0.01 | <0.001 |
| Observations |  | 11226 |  |  | 25303 |  |  | 12024 |  |  | 12024 |  |  | 29535 |  |
| R<sup>2</sup>/R<sup>2</sup> adjusted |  | 0.026 / 0.026 |  |  | 0.008 / 0.008 |  |  | 0.019 / 0.019 |  |  | 0.000 / 0.000 |  |  | 0.038 / 0.038 |  |

While we did not deliberately exclude certain habitat types, the data mainly consist of semi-natural to intensively managed grasslands and forests. Thus, the time series in ReSurveyGermany are biased with respect to habitat types. We assigned EUNIS habitat types to each plot record. This was accomplished by using the expert system EUNIS-ESy^78^ and the corresponding R code^79^. Plot records covered a total of 92 EUNIS habitat types out of the 150 ones distinghished for Germany. About 63% of the 23,641 plot records came from grasslands (level 1 EUNIS habitat R, n=14,849), followed by forests and other wooded lands (T, n= 5,440, 23%). In contrast, arable land (V, vegetated man-made habitats), which makes up more than 36% of the land cover in Germany, was only represented by 3% (816 plot records).

## Database organisation

A separate database was created for each project that contributed data, using the data-management software Turboveg 2^80^. The database is composed of two main tables, following the structure of Turboveg and common practice in vegetation science. The plot-species-abundance table contains six fields as described in Table 3. It is linked to the plot metadata (header file) through PROJECT_ID_RELEVE_NR, which is a unique Plot observation ID of a combination of PROJECT_ID (see Table 1) and the plot observation ID (called RELEVE_NR), the name of the observed taxon (TaxonName), the vertical layer (tree layer, shrub layer, herb layer, moss layer) in which the species was observed (LAYER) and the taxon’s cover in the plot (Cover_Perc). The latter was obtained by transforming the original cover classes in per cent cover. For example, the seven cover classes of theBraun-Blanquet scale, r, +, 1, 2, 3, 4, 5 were transformed to 1%, 2%, 3%, 13%, 38%, 63%, 88%, respectively. The other table is the so-called header file, which holds all important plot-level information, such as plot sizes, geographic location and vegetation structure for each plot observation ID (Table 4).

**Table 3:** Data structure of the Plot-species-abundance file of ReSurveyGermany. For Type: C = character, N = numeric, I = integer (n=23,641)

| Field name | Type | Description |
| --- | --- | --- |
| PROJECT_ID | I | Number of the resurvey project in ReSurveyGermany; see Table 1 |
| RELEVE_NR | I | Plot observation ID, only unique within a RS_PROJECT, usually the original plot observation ID from the underlying Turboveg 2 database |
| PROJECT_ID<br>_RELEVE_NR | C | Unique Plot observation ID, by which the project's plot-species-abundance file is linked to the header file, combination of PROJECT_ID and RELEVE_NR |
| LAYER | I | 0: No layer, 1: Tree layer (uppermost), 2: Tree layer (middle), 3: Tree layer (lowest), 4: Shrub layer (uppermost), 5: Shrub layer (low), 6: Herb layer, 7: Juveniles, 8: Seedling (< 1 year), 9: Moss layer. |
| TaxonName | C | Harmonized taxon name |
| Cover_Perc | C | Cover of the taxon in per cent |

**Table 4:** Data structure of the header file of ReSurveyGermany. For Type: C = character, N = numeric, I = integer. Per cent of NA values is given by dividing the number of NAs by (n=23,641)

| Field name | Type | Description | Number of NAs |
| --- | --- | --- | --- |
| RS_PROJECT | C | Unique name of the resurvey project; for the list of the 92 projects and the underlying references see Table 1 | 0 |
| PROJECT_ID | I | Number of the resurvey project in ReSurveyGermany; see Table 1 | 0 |
| RS_PLOT | C | Unique (within the site) code of the resurveyed plot; it is used to pair observations from different times recorded in the same plot; gives a unique identifier for the resurveyed plot or set of plots in time if combined with RS_PROJECT. Several plots in the same year might have the same RS_PLOT code if they have to be summarised for temporal comparisons. In these cases, they might also contain the community name. | 0 |
| RS_SITE | C | Name of the resurveyed site. For further details see LOCALITY. | 0 |
| LOCALITY | C | More detailed description of the locality of the resurveyed site (in German and if available) | 8,499 |
| RS_OBSERV | C | Unique code of the one-time observation; combination of RS_SITE, RS_PLOT, YEAR | 0 |
| RELEVE_NR | I | Plot observation ID, only unique within a RS_PROJECT, usually the original plot observation ID from the underlying Turboveg 2 database | 0 |
| PROJECT_ID<br>_RELEVE_NR | C | Unique Plot observation ID, by which the project's plot-species-abundance file is linked to the header file, combination of PROJECT_ID and RELEVE_NR | 0 |
| DATE | C | Date of the record (YYYYMMDD); the exact date if provided by the author, otherwise only the year and month or only year; if the year was not provided by the author, we took the year of the publication | 0 |
| YEAR | I | Year of the record (YYYY), extracted from DATE | 0 |
| SURF_AREA | N | Plot size [m<sup>2</sup>] (only stated if available) | 2064 |
| MANIPULATE | C | Binary information (Y/N) about whether the plot was part of a manipulative experiment ("Y") or not ("N"). If "Y", we chose the treatments representing the ambient land use. Observations with NA were to our knowledge not part of an experiment, and thus, can be treated as "N". | 15,020 |
| MANIPTYPE | C | Shows the type of treatment in the plot manipulation (partly in German and only if available). | 20,255 |
| LAND_USE | C | Land use, often identical with MANIPTYPE (mostly in German, also using the abbreviations used in the particular study, and only if available) | 18,149 |
| LOC_METHOD | C | Method of plot (re-)location, 1: Permanently marked plot isolated (i.e. somewhere within the site), 2: Marked plot in a grid (i.e. with regularly spaced neighbor plots), 3: Location with differential GPS, 4: Location with GPS, 5: Location from accurate map, 6: Location from a description, 7: Other | 12,607 |
| LOC_METH_COMMENT | C | Detailed description of the location method (if available) | 20,163 |
| LONGITUDE | N | Longitude of the plot in decimal degrees, coordinate system WGS-84; this coordinate should refer to the centre of the plot; coordinates were rounded to 2 digits of decimal degrees. | 0 |
| LATITUDE | N | Latitude of the plot in decimal degrees, rounded to 2 digits as LONGITUDE | 0 |
| PRECISION | I | Uncertainty in m, of coordinates for geographic position of plots, provided by the author or estimated if coordinates were taken from a topographic map. PRECISION refers to the true coordinates, not to those rounded to two digits. | 13,034 |
| GEO_LEV | C | Method of how the geographic location was obtained: GPS = Geographical positioning system, MTB = center of the German ordnance map, MTB_4 = center of a quadrant of the German ordnance map, POINT = all other | 0 |
| ALTITUDE | I | Elevation [m] (if available) | 14,723 |
| ASPECT | N | Compass direction of the slope in degrees [°], 0° = N, 90°=E etc. NA shows plot records either without aspect information or with aspect information when SLOPE is 0. | 16,572 |
| SLOPE | I | Inclination of the slope in degrees [°] | 18,962 |
| COUNTRY | C | DE for Germany | 0 |
| EUNIS |  | EUNIS level 3 code of the habitat, as obtained by applying the expert system EUNIS-ESy<sup>78</sup> and the corresponding R code<sup>79</sup>. | 0 |
| COVERSCALE | C | Cover scale used for the plot record. 00 = no scale, cover estimated in per cent (%), 01 = Braun/Blanquet (old), 02 = Braun/Blanquet (new), 03 = Londo, 04 = Presence/Absence, 10 = Reichelt & Wilmanns 1973 (short), 26 = Londo (short), 29 = Londo per cent classes, 30 = Londo (modified, in project 9, Sperle et al. unpublished), 31 = Maas & Kohler 1983 (in project 86, Poschlod et al. 2010), 50 = Londo (modified, in project 89, Bruelheide & Luginbühl 2009) | 0 |
| REFERENCE | C | Reference number in GVRD, 6 digits referring to the bibliographic reference, found in ReSurveyGermanyReference.csv | 51 |
| YEAR_PUBL | I | Year of the publication (if available) | 18,057 |
| TABLE_NR | C | Number of the table in the original publication | 12,659 |
| TABNAME |  | Name of the table in the original publication | 8,402 |
| NR_IN_TAB | C | Column name in the TABLE_NR | 3,789 |
| ORIG_NR |  | Name of the plot given by the author in the original publication | 10,172 |
| ORIG_DB |  | Name of original Turboveg file, to be used internally for backtracking changes | 19,700 |
| COV_TOTAL | I | Total cover of all layers [%] (if available) | 18,704 |
| COV_TREES | I | Cover of the tree layer [%] (if available) | 20,554 |
| COV_SHRUB<br>S | I | Cover of the shrub layer [%] (if available) | 20,520 |
| COV_HERBS | I | Cover of the herb layer [%] (if available) | 11,964 |
| COV_MOSSE<br>S | I | Cover of the moss layer (bryophytes and lichens) [%] (if available) | 17,512 |
| COV_LITTER | I | Cover of the litter layer on the ground [%] (if available) | 20,786 |
| COV_ROCK | I | Cover of the rocks on the plot surface [%] (if available) | 21,697 |
| TREE_HIGH | I | Height of the upper tree layer [m] (if available) | 22,317 |
| TREE_LOW | I | Height of the lower tree layer [m] (if available) | 23,107 |
| SHRUB_HIG<br>H | N | Height of the upper shrub layer [m] (if available) | 22,848 |
| SHRUB_LOW | N | Height of the lower shrub layer [m] (if available) | 23,478 |
| HERB_HIGH |  | Mean height of the upper herb layer [cm] (if available) | 20,311 |
| HERB_LOW |  | Mean height of the lower herb layer [cm] (if available) | 22,317 |
| HERB_MAX |  | Maximum height of the herb layer [cm] (if available) | 22,627 |

The taxon names in the plot-species-abundance table were standardised using German SL 1.3^81^. The nomenclature for vascular plants followed Wisskirchen et al.^82^, with additional aggregations to higher taxonomic levels according to German SL 1.3. As some authors recorded subspecies and other infraspecific taxa, species were aggregated at the species level, using vegdata^83^. Some closely related species that, from our experience, are often mistaken in the field were merged at the aggregate or genus level. Species aggregates were also used when different taxon names of the same aggregate occurred in different projects, to prevent that the same taxon might appear under different taxon names. We used our own R code to merge taxon names and the notation of the ESy expert system^78^ to protocol all steps. The species harmonisation forms the first section of the ESy system and shows which taxon names were aggregated under the name of a broader taxonomic concept (Table 5). In addition, within single projects, we used customised aggregations and segregations when the same taxa were reported with different taxonomic levels at different points in time in the same plot resurvey IDs (Table 6). For example, in all years of a time series of a specific plot *Orchis militaris* was reported but in one year *Orchis* spec. was recorded at the genus level. Unaccounted for, such a leap between taxonomic levels within a time series would result in incorrect species change observations. To avoid losing the predominating information at the species level by aggregating all records to *Orchis*, we assumed that the taxon was also *Orchis militaris* in the particular year when only the genus level was reported. If more than one taxon occurred in previous years, we equally distributed the cover values among those taxa. This happened for example when a record was taken late in spring when the two species *Anemone nemorosa* and *A. ranunculoides* could no longer be distinguished.

**Table 5:** List of all taxa that were harmonised across all projects. The format of the list follows the rules of the ESy system^78^. The taxon names that were aggregated below a broader concept name are indented using five blanks. The number to the right shows the German SL 1.3^81^ number for each taxon.

|  |  |
| --- | --- |
| Achillea atrata agg. | 18 |
| Achillea atrata | 19 |
| Achillea atrata agg. | 18 |
| Achillea millefolium agg. | 27 |
| Achillea millefolium | 31 |
| Achillea millefolium agg. | 27 |
| Achillea millefolium subsp. collina | 20096 |
| Achillea millefolium subsp. millefolium | 32 |
| Achillea pannonica | 34 |
| Achillea setacea | 36 |
| Acinos arvensis | 49 |
| Acinos arvensis | 49 |
| Calamintha acinos | 976 |
| Satureja acinos | 23760 |
| Aconitum lycoctonum | 14242 |
| Aconitum lycoctonum | 14242 |
| Aconitum lycoctonum subsp. vulparia | 20209 |
| Aconitum vulparia | 68 |
| Adonis aestivalis | 76 |
| Adonis aestivalis | 76 |
| Adonis aestivalis var. citrinus | 27247 |
| Agrimonia eupatoria | 99 |
| Agrimonia eupatoria | 99 |
| Agrimonia eupatoria subsp. eupatoria | 100 |
| Agrostis canina agg. | 120 |
| Agrostis canina | 121 |
| Agrostis coarctata | 20180 |
| Agrostis stricta | 122 |
| Agrostis vinealis | 20684 |
| Agrostis capillaris | 20178 |
| Agrostis capillaris | 20178 |
| Agrostis capillaris subsp. oreophila | 7086 |
| Agrostis tenuis | 130 |
| Agrostis vulgaris | 20179 |
| Agrostis stolonifera agg. | 127 |
| Agrostis alba var. stolonifera | 27181 |
| Agrostis gigantea | 128 |
| Agrostis stolonifera | 129 |
| Agrostis stolonifera agg. | 127 |
| Agrostis stolonifera subsp. stolonifera | 6550 |
| Aira caryophyllea | 26920 |
| Aira caryophyllea | 26920 |
| Aira caryophyllea subsp. caryophyllea | 20218 |
| Alchemilla conjuncta agg. | 154 |
| Alchemilla conjuncta agg. | 154 |
| Alchemilla hoppeana | 166 |
| Alchemilla nitida | 10027 |
| Alchemilla fissa agg. | 177 |
| Alchemilla fallax | 179 |
| Alchemilla fissa | 180 |
| Alchemilla fissa agg. | 177 |
| Alchemilla hybrida agg. | 190 |
| Alchemilla flabellata | 194 |
| Alchemilla glaucescens | 195 |
| Alchemilla hybrida agg. | 190 |
| Alchemilla mollis | 200 |
| *Alchemilla acutiloba* var. *mollis* | 20237 |
| *Alchemilla vulgaris* agg. | 209 |
| *Alchemilla acutiloba* var. *stellata* | 11694 |
| *Alchemilla crinita* | 219 |
| *Alchemilla glabra* | 235 |
| *Alchemilla micans* | 20278 |
| *Alchemilla monticola* | 245 |
| *Alchemilla subcrenata* | 265 |
| *Alchemilla vulgaris* | 26467 |
| *Alchemilla vulgaris* agg. | 209 |
| *Alchemilla vulgaris* auct. | 20289 |
| *Alchemilla xanthochlora* | 273 |
| *Alisma plantago-aquatica* agg. | 275 |
| *Alisma gramineum* | 276 |
| *Alisma lanceolatum* | 277 |
| *Alisma plantago-aquatica* | 278 |
| *Alisma plantago-aquatica* agg. | 275 |
| *Allium scorodoprasum* | 12756 |
| *Allium rotundum* | 310 |
| *Allium scorodoprasum* | 12756 |
| *Allium scorodoprasum* subsp. *rotundum* | 20326 |
| *Allium senescens* | 6583 |
| *Allium montanum* | 294 |
| *Allium senescens* | 6583 |
| *Allium senescens* subsp. *montanum* | 20321 |
| *Allium senescens* var. *montanum* | 12712 |
| *Alopecurus pratensis* agg. | 334 |
| *Alopecurus pratensis* | 336 |
| *Alopecurus pratensis* agg. | 334 |
| *Alyssum alyssoides* | 345 |
| *Alyssum alyssoides* | 345 |
| *Alyssum calycinum* | 20351 |
| *Alyssum montanum* | 350 |
| *Alyssum montanum* | 350 |
| *Alyssum montanum* subsp. *gmelinii* | 26457 |
| *Alyssum montanum* subsp. *montanum* | 26456 |
| *Anagallis arvensis* | 393 |
| *Anagallis arvensis* | 393 |
| *Anagallis arvensis* subsp. *arvensis* | 394 |
| *Anemone narcissiflora* | 434 |
| *Anemonastrum narcissiflorum* | 20423 |
| *Anemone narcissiflora* | 434 |
| *Anthoxanthum aristatum* | 20071 |
| *Anthoxanthum aristatum* | 20071 |
| *Anthoxanthum puelii* | 466 |
| *Anthoxanthum odoratum* agg. | 463 |
| *Anthoxanthum alpinum* | 464 |
| *Anthoxanthum odoratum* | 465 |
| *Anthoxanthum odoratum* agg. | 463 |
| *Anthriscus sylvestris* agg. | 469 |
| *Anthriscus sylvestris* | 473 |
| *Anthriscus sylvestris* agg. | 469 |
| *Anthriscus sylvestris* subsp. *alpestris* | 20460 |
| *Anthriscus sylvestris* subsp. *sylvestris* | 20463 |
| *Anthyllis vulneraria* | 477 |
| *Anthyllis vulneraria* | 477 |
| *Anthyllis vulneraria* subsp. *carpatica* | 479 |
| *Anthyllis vulneraria* subsp. *pseudovulneraria* | 14783 |
| *Aphanes inexpectata* | 494 |
| *Aphanes inexpectata* | 494 |
| *Aphanes microcarpa* | 6554 |
| *Aquilegia vulgaris* agg. | 507 |
| *Aquilegia atrata* | 508 |
| *Aquilegia vulgaris* | 510 |
| *Arabis bellidifolia* | 20511 |
| *Arabis bellidifolia* subsp. *bellidifolia* | 20512 |
| *Arabis glabra* | 520 |
| *Arabis glabra* | 520 |
| *Turritis glabra* | 24405 |
| *Arabis hirsuta* agg. | 521 |
| *Arabis hirsuta* | 523 |
| *Arabis nemorensis* | 20072 |
| *Arctium minus* | 13702 |
| *Arctium minus* | 13702 |
| *Arctium minus* agg. | 547 |
| *Lappa communis* | 14348 |
| *Arenaria serpyllifolia* agg. | 563 |
| *Arenaria serpyllifolia* | 13703 |
| *Arenaria serpyllifolia* agg. | 563 |
| *Armeria maritima* | 581 |
| *Armeria elongata* | 577 |
| *Armeria maritima* | 581 |
| *Armeria maritima* agg. | 575 |
| *Armeria maritima* subsp. *elongata* | 20584 |
| *Armeria maritima* var. *elongata* | 20590 |
| *Armeria vulgaris* | 26026 |
| *Artemisia campestris* agg. | 594 |
| *Artemisia campestris* | 596 |
| *Artemisia campestris* agg. | 594 |
| *Artemisia campestris* subsp. *campestris* | 6466 |
| *Artemisia vulgaris* agg. | 616 |
| *Artemisia vulgaris* | 618 |
| *Artemisia vulgaris* agg. | 616 |
| *Arum maculatum* agg. | 622 |
| *Arum maculatum* | 625 |
| *Arum maculatum* agg. | 622 |
| *Asplenium ruta-muraria* | 674 |
| *Asplenium ruta-muraria* | 674 |
| *Asplenium ruta-muraria* subsp. *ruta-muraria* | 676 |
| *Aster lanceolatus* agg. | 15106 |
| *Aster lanceolatus* | 693 |
| *Aster parviflorus* | 15085 |
| *Atriplex prostrata* agg. | 20706 |
| *Atriplex calotheca* | 756 |
| *Atriplex hastata* | 20693 |
| *Atriplex hastata* agg. | 755 |
| *Atriplex latifolia* | 760 |
| *Atriplex prostrata* | 762 |
| *Atriplex sagittata* | 754 |
| *Atriplex nitens* | 20702 |
| *Aurinia saxatilis* | 777 |
| *Alyssum saxatile* | 354 |
| *Betonica officinalis* | 826 |
| *Betonica officinalis* | 826 |
| *Stachys officinalis* | 24165 |
| *Betula alba* agg. | 99014 |
| *Betula pendula* | 829 |
| *Betula verrucosa* | 20786 |
| *Betula pubescens* | 830 |
| *Betula pubescens* | 830 |
| *Betula pubescens* subsp. *carpatica* | 6472 |
| *Betula pubescens* subsp. *pubescens* | 6471 |
| *Bidens frondosa* | 834 |
| *Bidens frondosa* | 834 |
| *Bidens melanocarpa* | 20788 |
| *Bidens tripartita* | 836 |
| Bidens tripartita | 836 |
| Bidens tripartita subsp. tripartita | 838 |
| Biscutella laevigata | 844 |
| Biscutella laevigata | 844 |
| Biscutella laevigata subsp. kernerii | 6652 |
| Bistorta officinalis | 27744 |
| Bistorta officinalis | 27744 |
| Persicaria bistorta | 23029 |
| Polygonum bistorta | 4420 |
| Bistorta vivipara | 20801 |
| Bistorta vivipara | 20801 |
| Persicaria vivipara | 23045 |
| Polygonum viviparum | 4437 |
| Bolboschoenus maritimus | 852 |
| Bolboschoenus maritimus | 852 |
| Schoenoplectus maritimus | 23820 |
| Bothriochloa ischaemum | 854 |
| Andropogon ischaemum | 20417 |
| Bothriochloa ischaemum | 854 |
| Dichanthium ischaemum | 21538 |
| Brachypodium pinnatum agg. | 862 |
| Brachypodium pinnatum | 863 |
| Brachypodium pinnatum agg. | 862 |
| Brachypodium rupestre | 864 |
| Bromus arvensis | 883 |
| Bromus arvensis | 883 |
| Bromus arvensis subsp. arvensis | 7191 |
| Bromus hordeaceus agg. | 895 |
| Bromus hordeaceus | 896 |
| Bromus hordeaceus agg. | 895 |
| Bromus hordeaceus subsp. hordeaceus | 897 |
| Bromus mollis | 901 |
| Bromus racemosus agg. | 908 |
| Bromus commutatus | 909 |
| Bromus racemosus | 910 |
| Bromus ramosus agg. | 911 |
| Bromus benekenii | 912 |
| Bromus ramosus | 913 |
| Bromus ramosus agg. | 911 |
| Callitriche palustris agg. | 999 |
| Callitriche obtusangula | 1003 |
| Callitriche palustris | 1004 |
| Callitriche palustris agg. | 999 |
| Callitriche stagnalis | 1006 |
| Campanula rotundifolia agg. | 1057 |
| Campanula rotundifolia | 1072 |
| Campanula rotundifolia agg. | 1057 |
| Campanula scheuchzeri | 1073 |
| Cardamine bulbifera | 20944 |
| Cardamine bulbifera | 20944 |
| Dentaria bulbifera | 1896 |
| Cardamine heptaphylla | 20951 |
| Dentaria heptaphyllos | 91015 |
| Cardamine pratensis agg. | 1105 |
| Cardamine dentata | 20945 |
| Cardamine palustris | 1109 |
| Cardamine pratensis | 15133 |
| Cardamine pratensis agg. | 1105 |
| Cardaminopsis arenosa | 1114 |
| Arabis arenosa | 20509 |
| Cardaminopsis arenosa | 1114 |
| Cardaminopsis halleri | 1116 |
| Arabis halleri | 20518 |
| Cardaminopsis halleri | 1116 |
| Cardaminopsis petraea | 1117 |
| Arabis hispida | 20526 |
| Carduus nutans agg. | 1140 |
| Carduus nutans | 1143 |
| Carduus nutans agg. | 1140 |
| Carex acuta agg. | 26775 |
| Carex acuta | 20989 |
| Carex acuta subsp. tricostata | 6685 |
| Carex gracilis | 1230 |
| Carex arenaria agg. | 1159 |
| Carex arenaria | 1160 |
| Carex ligerica | 1161 |
| Carex pseudobrizoides | 1162 |
| Carex atrata agg. | 1164 |
| Carex atrata | 14018 |
| Carex atrata agg. | 1164 |
| Carex elata | 1204 |
| Carex elata | 1204 |
| Carex elata subsp. elata | 1205 |
| Carex flacca | 1216 |
| Carex flacca | 1216 |
| Carex glauca | 21026 |
| Carex flava agg. | 1219 |
| Carex demissa | 20011 |
| Carex flava | 1220 |
| Carex flava agg. | 1219 |
| Carex flava var. oederi | 25010 |
| Carex lepidocarpa | 1222 |
| Carex oederi | 1223 |
| Carex serotina | 21070 |
| Carex tumidicarpa | 1225 |
| Carex viridula | 6693 |
| Carex muricata agg. | 1249 |
| Carex divulsa | 1250 |
| Carex leersiana | 1251 |
| Carex muricata | 6687 |
| Carex muricata agg. | 1249 |
| Carex muricata subsp. muricata | 6689 |
| Carex pairae | 1252 |
| Carex spicata | 1253 |
| Carex nigra agg. | 1254 |
| Carex fusca | 21024 |
| Carex nigra | 1256 |
| Carex nigra agg. | 1254 |
| Carex ornithopoda agg. | 1262 |
| Carex ornithopoda | 1263 |
| Carex ornithopoda agg. | 1262 |
| Carex ovalis | 21055 |
| Carex leporina | 1240 |
| Carex ovalis | 21055 |
| Carex vulpina agg. | 1301 |
| Carex otrubae | 1302 |
| Carex vulpina | 1303 |
| Carex vulpina agg. | 1301 |
| Carex × elytroides | 10122 |
| Carex × elytroides | 10122 |
| Carex acuta × nigra | 90526 |
| Carex × figertii | 90596 |
| Carex davalliana × dioica | 90549 |
| Carex × xanthocarpa | 10137 |
| Carex flava × hostiana | 90562 |
| Carlina acaulis | 1306 |
| *Carlina acaulis* | 1306 |
| *Carlina acaulis* subsp. *simplex* | 1308 |
| *Carlina vulgaris* | 1312 |
| *Carlina vulgaris* | 1315 |
| *Carlina vulgaris* subsp. *vulgaris* | 21094 |
| *Castanea sativa* | 1323 |
| *Castanea sativa* | 1323 |
| *Castanea vesca* | 21097 |
| *Caucalis platycarpos* | 1329 |
| *Caucalis lappula* | 21108 |
| *Caucalis platycarpos* | 1329 |
| *Centaurea jacea* | 1347 |
| *Centaurea angustifolia* | 21122 |
| *Centaurea jacea* | 1347 |
| *Centaurea jacea* agg. | 24981 |
| *Centaurea jacea* subsp. *amara* | 21132 |
| *Centaurea jacea* subsp. *angustifolia* | 1348 |
| *Centaurea nigra* | 26577 |
| *Centaurea nemoralis* | 21138 |
| *Centaurea nigra* | 26577 |
| *Centaurea nigra* subsp. *nemoralis* | 1370 |
| *Centaurea scabiosa* | 1390 |
| *Centaurea scabiosa* | 1390 |
| *Centaurea scabiosa* subsp. *scabiosa* | 1397 |
| *Centaurea stoebe* | 25004 |
| *Centaurea maculosa* subsp. *rhenana* | 21134 |
| *Centaurea rhenana* | 21146 |
| *Centaurea stoebe* | 25004 |
| *Centaureum erythraea* | 1406 |
| *Centaureum erythraea* | 1406 |
| *Centaureum erythraea* subsp. *erythraea* | 1407 |
| *Centaureum umbellatum* | 21156 |
| *Centaureum littorale* | 1409 |
| *Centaureum littorale* | 1409 |
| *Centaureum minus* | 7104 |
| *Cerastium arvense* | 1431 |
| *Cerastium arvense* | 1431 |
| *Cerastium arvense* subsp. *arvense* | 1432 |
| *Cerastium fontanum* agg. | 1449 |
| *Cerastium caespitosum* | 21178 |
| *Cerastium fontanum* | 1450 |
| *Cerastium fontanum* agg. | 1449 |
| *Cerastium fontanum* subsp. *triviale* | 21187 |
| *Cerastium holosteoides* | 1451 |
| *Cerastium triviale* | 25269 |
| *Cerastium vulgare* | 21213 |
| *Cerastium pumilum* agg. | 1460 |
| *Cerastium glutinosum* | 1461 |
| *Cerastium pallens* | 21198 |
| *Cerastium pumilum* | 1462 |
| *Cerastium pumilum* agg. | 1460 |
| *Ceratocarpus claviculata* | 21221 |
| *Ceratocarpus claviculata* | 21221 |
| *Corydalis claviculata* | 1670 |
| *Cerinth glabra* | 1473 |
| *Cerinth alpina* | 21227 |
| *Chaerophyllum hirsutum* | 26947 |
| *Chaerophyllum hirsutum* | 26947 |
| *Chaerophyllum hirsutum* agg. | 1490 |
| *Chaerophyllum villarsii* | 1493 |
| *Chamaecytisus ratisbonensis* | 1504 |
| *Chamaecytisus ratisbonensis* | 1504 |
| *Cytisus ratisbonensis* | 21484 |
| Chamaecytisus supinus | 1506 |
| Chamaecytisus supinus | 1506 |
| Cytisus supinus | 21486 |
| Chamaespartium sagittale | 1509 |
| Chamaespartium sagittale | 1509 |
| Genista sagittalis | 21992 |
| Genistella sagittalis | 2615 |
| Chenopodium album agg. | 1514 |
| Chenopodium album | 1515 |
| Chenopodium album agg. | 1514 |
| Chenopodium strictum subsp. striatifforme | 6482 |
| Cirsium acaule | 1556 |
| Cirsium acaule | 1556 |
| Cirsium acaulon | 21327 |
| Cirsium heterophyllum | 1569 |
| Cirsium helenioides | 21329 |
| Cirsium heterophyllum | 1569 |
| Cirsium vulgare | 1579 |
| Cirsium lanceolatum | 21330 |
| Cirsium vulgare | 1579 |
| Cirsium × rigens | 26068 |
| Cirsium acaule × oleraceum | 90817 |
| Clinopodium vulgare | 1593 |
| Calamintha clinopodium | 978 |
| Consolida regalis | 1627 |
| Consolida regalis | 1627 |
| Delphinium consolida | 21510 |
| Convolvulus | 60816 |
| Convolvulus | 60816 |
| Convolvulus arvensis | 1632 |
| Conyza canadensis | 1638 |
| Conyza canadensis | 1638 |
| Erigeron canadensis | 21687 |
| Crataegus | 61165 |
| Crataegus curvisepala | 1697 |
| Crataegus laevigata agg. | 1701 |
| Crataegus laevigata | 1701 |
| Crataegus lindmanii | 1699 |
| Crataegus monogyna | 1707 |
| Crataegus monogyna subsp. monogyna | 1708 |
| Crataegus monogyna subsp. nordica | 1709 |
| Crataegus monogyna var. monogyna | 90937 |
| Crataegus oxyacantha | 21400 |
| Crataegus rhipidophylla | 26677 |
| Crataegus × macrocarpa | 1705 |
| Crataegus × media | 50021 |
| Crataegus calycina | 6733 |
| Crepis bocconi | 21408 |
| Crepis pontana | 1735 |
| Cruciata laevipes | 1766 |
| Cruciata laevipes | 1766 |
| Galium cruciata | 21963 |
| Cystopteris fragilis agg. | 1825 |
| Cystopteris alpina | 21473 |
| Cystopteris fragilis | 1827 |
| Cytisus nigricans | 21483 |
| Cytisus nigricans | 21483 |
| Lembotropis nigricans | 3333 |
| Cytisus scoparius | 1837 |
| Cytisus scoparius | 1837 |
| Sarothamnus scoparius | 5245 |
| Dactylis glomerata agg. | 1842 |
| Dactylis | 397 |
| Dactylis aschersoniana | 21487 |
| Dactylis glomerata | 1843 |
| Dactylis glomerata agg. | 1842 |
| Dactylis glomerata subsp. glomerata | 13464 |
| Dactylis polygama | 1846 |
| Dactylorhiza maculata agg. | 1852 |
| Dactylorhiza fuchsii | 1853 |
| Dactylorhiza fuchsii × maculata | 90986 |
| Dactylorhiza maculata | 1857 |
| Dactylorhiza maculata agg. | 1852 |
| Dactylorhiza majalis agg. | 1861 |
| Dactylorhiza majalis | 1862 |
| Dactylorhiza majalis agg. | 1861 |
| Dactylorhiza majalis subsp. majalis | 1866 |
| Dactylorhiza traunsteineri | 1871 |
| Dactylorhiza × carnea | 13760 |
| Dactylorhiza maculata × incarnata | 90995 |
| Danthonia decumbens | 1874 |
| Danthonia decumbens | 1874 |
| Danthonia decumbens subsp. decumbens | 1876 |
| Sieglingia decumbens | 24019 |
| Daucus carota | 1886 |
| Daucus | 61271 |
| Daucus carota | 1886 |
| Daucus carota subsp. carota | 6753 |
| Deschampsia cespitosa agg. | 1903 |
| Deschampsia cespitosa | 1904 |
| Deschampsia cespitosa agg. | 1903 |
| Deschampsia flexuosa | 20725 |
| Aira flexuosa | 20226 |
| Avenella flexuosa | 783 |
| Deschampsia flexuosa | 20725 |
| Dianthus gratianopolitanus | 1934 |
| Dianthus caesius | 21522 |
| Dianthus gratianopolitanus | 1934 |
| Dianthus superbus | 1949 |
| Dianthus superbus | 1949 |
| Dianthus superbus subsp. superbus | 1952 |
| Digitalis grandiflora | 1960 |
| Digitalis ambigua | 21540 |
| Dipsacus fullonum | 1980 |
| Dipsacus fullonum | 1980 |
| Dipsacus sylvestris | 20021 |
| Draba aizoides agg. | 2000 |
| Draba aizoides | 2001 |
| Draba aizoides agg. | 2000 |
| Drosera longifolia | 21588 |
| Drosera anglica | 2025 |
| Drosera longifolia | 21588 |
| Dryopteris carthusiana agg. | 2030 |
| Dryopteris × deweveri | 50029 |
| Dryopteris carthusiana | 2032 |
| Dryopteris carthusiana × dilatata | 91118 |
| Dryopteris carthusiana agg. | 2030 |
| Dryopteris dilatata | 2033 |
| Dryopteris expansa | 2031 |
| Dryopteris spinulosa | 21604 |
| Dryopteris filix-mas agg. s. l. | 94728 |
| Dryopteris affinis | 2038 |
| Dryopteris filix-mas | 2037 |
| Dryopteris filix-mas agg. | 2035 |
| Eleocharis palustris agg. | 2087 |
| Eleocharis palustris | 2091 |
| Eleocharis palustris agg. | 2087 |
| Eleocharis uniglumis | 2094 |
| Scirpus palustris | 23856 |
| Elymus arenosus | 27779 |
| Elymus repens subsp. arenosus | 6543 |
| Elymus athericus | 27782 |
| Elymus pungens | 27905 |
| Elymus caninus | 20145 |
| Agropyron caninum | 103 |
| Elymus caninus | 20145 |
| Roegneria canina | 23598 |
| Elymus repens | 27778 |
| Agropyron repens | 27914 |
| Agropyron repens subsp. caesium | 6541 |
| Elymus repens | 27778 |
| Elymus repens subsp. repens | 27781 |
| Elytrigia repens | 21639 |
| Triticum repens | 24393 |
| Empetrum nigrum agg. | 2103 |
| Empetrum nigrum | 2105 |
| Empetrum nigrum agg. | 2103 |
| Epilobium angustifolium | 2113 |
| Chamaenerion angustifolium | 21237 |
| Epilobium angustifolium | 2113 |
| Epilobium ciliatum | 21642 |
| Epilobium adenocaulon | 2109 |
| Epilobium ciliatum | 21642 |
| Epilobium tetragonum | 2126 |
| Epilobium tetragonum | 2126 |
| Epilobium tetragonum subsp. lamyi | 2127 |
| Epilobium tetragonum subsp. tetragonum | 2128 |
| Epipactis atrorubens | 2130 |
| Epipactis atropurpurea | 21653 |
| Epipactis atrorubens | 2130 |
| Epipactis rubiginosa | 21664 |
| Epipactis helleborine agg. | 2131 |
| Epipactis helleborine | 2134 |
| Epipactis latifolia | 21660 |
| Equisetum fluviatile | 2143 |
| Equisetum fluviatile | 2143 |
| Equisetum limosum | 21674 |
| Equisetum × mildeanum | 91305 |
| Equisetum pratense × sylvaticum | 91302 |
| Erica carnea | 21680 |
| Erica carnea | 21680 |
| Erica herbacea | 2163 |
| Erigeron acris | 2167 |
| Erigeron acris | 2167 |
| Erigeron acris subsp. acris | 2168 |
| Erigeron annuus | 2178 |
| Erigeron annuus | 2178 |
| Erigeron strigosus | 21695 |
| Erigeron glabratus | 21689 |
| Erigeron glabratus | 21689 |
| Erigeron polymorphus | 2186 |
| Erodium cicutarium agg. | 2197 |
| Erodium cicutarium | 2199 |
| Erodium cicutarium agg. | 2197 |
| Erophila verna | 12356 |
| Draba verna | 21586 |
| Erophila verna | 12356 |
| Erophila verna agg. | 2205 |
| Erophila verna subsp. verna | 21715 |
| *Erysimum maschallianum* | 2229 |
| *Erysimum durum* | 21726 |
| *Erysimum odoratum* | 2234 |
| *Erysimum erysimoides* | 21727 |
| *Erysimum odoratum* | 2234 |
| *Euphorbia verrucosa* | 2309 |
| *Euphorbia brittingeri* | 21740 |
| *Euphorbia verrucosa* | 2309 |
| *Euphrasia officinalis* | 13816 |
| *Euphrasia officinalis* | 13816 |
| *Euphrasia officinalis* subsp. *rostkoviana* | 26254 |
| *Euphrasia rostkoviana* | 2332 |
| *Euphrasia rostkoviana* agg. | 2329 |
| *Fallopia convolvulus* | 2359 |
| *Fallopia convolvulus* | 2359 |
| *Polygonum convolvulus* | 23228 |
| *Festuca brevipila* | 26591 |
| *Festuca brevipila* | 26591 |
| *Festuca duriuscula* | 21820 |
| *Festuca trachyphylla* | 2406 |
| *Festuca filiformis* | 21824 |
| *Festuca capillata* | 21813 |
| *Festuca filiformis* | 21824 |
| *Festuca tenuifolia* | 2405 |
| *Festuca ovina* agg. | 2385 |
| *Festuca guestfalica* et *ovina* | 91404 |
| *Festuca guestfalica* | 6483 |
| *Festuca lemanii* | 94730 |
| *Festuca ovina* | 2396 |
| *Festuca ovina* s. l. | 91415 |
| *Festuca ovina* subsp. *ovina* | 91418 |
| *Festuca ovina* var. *firmula* | 21845 |
| *Festuca ovina* agg. | 2385 |
| *Festuca ovina* subsp. *glauca* | 10352 |
| *Festuca pallens* | 2397 |
| *Festuca cinerea* | 2387 |
| *Festuca glauca* | 2392 |
| *Festuca glaucina* | 91400 |
| *Festuca pallens* | 2397 |
| *Festuca pallens* * *glaucina* | 91420 |
| *Festuca pallens* subsp. *pallens* | 6792 |
| *Festuca quadriflora* | 21852 |
| *Festuca pumila* | 2416 |
| *Festuca quadriflora* | 21852 |
| *Festuca rubra* agg. | 2417 |
| *Festuca heteromalla* | 7346 |
| *Festuca nigrescens* | 2420 |
| *Festuca nigrescens* subsp. *nigrescens* | 91414 |
| *Festuca rubra* | 2421 |
| *Festuca rubra* agg. | 2417 |
| *Festuca rubra* subsp. *commutata* | 21856 |
| *Festuca rubra* subsp. *fallax* | 13510 |
| *Festuca rubra* subsp. *rubra* | 2425 |
| *Festuca rubra* var. *genuina* | 21863 |
| *Festuca trichophylla* | 2426 |
| *Festuca rupicola* | 2402 |
| *Festuca rupicola* | 2402 |
| *Festuca sulcata* | 27166 |
| *Festuca valesiaca* subsp. *sulcata* | 10354 |
| *Festuca valesiaca* | 13712 |
| *Festuca pulchra* | 13523 |
| *Festuca valesiaca* | 13712 |
| *Festuca violacea* agg. | 2441 |
| Festuca norica | 2443 |
| Festuca violacea agg. | 2441 |
| Filipendula vulgaris | 2462 |
| Filipendula hexapetala | 21900 |
| Filipendula vulgaris | 2462 |
| Frangula alnus | 2472 |
| Frangula alnus | 2472 |
| Rhamnus frangula | 23535 |
| Fumana procumbens | 2486 |
| Fumana procumbens | 2486 |
| Fumana vulgaris | 21921 |
| Galeopsis ladanum agg. | 2520 |
| Galeopsis angustifolia | 2521 |
| Galeopsis ladanum | 2522 |
| Galeopsis tetrahit agg. | 2526 |
| Galeopsis bifida | 2527 |
| Galeopsis tetrahit | 2528 |
| Galeopsis tetrahit agg. | 2526 |
| Galium aparine agg. | 2532 |
| Galium aparine | 2533 |
| Galium aparine agg. | 2532 |
| Galium spurium | 2534 |
| Galium glaucum | 2542 |
| Asperula glauca | 20637 |
| Galium glaucum | 2542 |
| Galium mollugo agg. | 2548 |
| Galium album | 2549 |
| Galium album subsp. album | 2550 |
| Galium mollugo | 2555 |
| Galium mollugo agg. | 2548 |
| Galium palustre | 2564 |
| Galium elongatum | 2563 |
| Galium palustre | 2564 |
| Galium palustre agg. | 2561 |
| Galium palustre subsp. elongatum | 21974 |
| Galium palustre subsp. palustre | 21975 |
| Galium pusillum agg. | 2569 |
| Galium anisophyllum | 2570 |
| Galium pumilum | 2572 |
| Galium pusillum agg. | 2569 |
| Galium valdepilosum | 2577 |
| Galium rotundifolium | 2579 |
| Galium scabrum | 21982 |
| Galium saxatile | 2585 |
| Galium hircynicum | 2543 |
| Galium saxatile | 2585 |
| Galium verum agg. s. l. | 94733 |
| Galium x pomeranicum | 2599 |
| Galium verum | 2600 |
| Galium verum agg. | 2598 |
| Galium verum subsp. verum | 24929 |
| Galium wirtgenii | 2601 |
| Gentiana acaulis agg. | 91530 |
| Gentiana acaulis | 2616 |
| Gentiana clusii | 2624 |
| Gentianella ciliata | 2656 |
| Gentiana ciliata | 22041 |
| Gentianella ciliata | 2656 |
| Gentianopsis ciliata | 22040 |
| Gentianella germanica agg. | 2657 |
| Gentiana germanica | 22044 |
| Gentianella aspera | 2659 |
| Gentianella germanica | 2662 |
| Gentianella germanica agg. | 2657 |
| Gentianella germanica subsp. germanica | 7352 |
| Gentianella lutescens | 2665 |
| Geum × intermedium | 50040 |
| Geum rivale × urbanum | 91547 |
| Glechoma hederacea agg. | 2710 |
| Glechoma hederacea | 2711 |
| Glechoma hederacea agg. | 2710 |
| Globularia punctata | 2718 |
| Globularia bisnagarica | 2718 |
| Globularia elongata | 22079 |
| Globularia punctata | 2718 |
| Globularia vulgaris | 26117 |
| Globularia willkommii | 22080 |
| Glyceria fluitans agg. | 2719 |
| Glyceria declinata | 2720 |
| Glyceria fluitans | 2721 |
| Glyceria fluitans agg. | 2719 |
| Glyceria notata | 26260 |
| Helianthemum nummularium | 26596 |
| Helianthemum nummularium | 26596 |
| Helianthemum nummularium agg. | 2774 |
| Helianthemum nummularium subsp. obscurum | 22127 |
| Helianthemum nummularium subsp. ovatum | 22128 |
| Helianthemum ovatum | 2780 |
| Helictotrichon pratense | 20728 |
| Avena pratensis | 20717 |
| Avenochloa pratensis | 788 |
| Avenula pratensis | 20732 |
| Helictotrichon pratense | 20728 |
| Helictotrichon pubescens | 20729 |
| Avena pubescens | 20718 |
| Avenochloa pubescens | 789 |
| Avenula pubescens | 20733 |
| Helictotrichon pubescens | 20729 |
| Helictotrichon versicolor | 20730 |
| Avenula versicolor | 20734 |
| Hepatica nobilis | 2815 |
| Anemone hepatica | 20422 |
| Hepatica nobilis | 2815 |
| Hepatica triloba | 10391 |
| Herniaria glabra | 2832 |
| Herniaria glabra | 2832 |
| Herniaria glabra subsp. glabra | 2833 |
| Hieracium angustifolium | 25621 |
| Hieracium glaciale | 2885 |
| Hieracium caespitosum | 2866 |
| Pilosella caespitosa | 12173 |
| Hieracium lactucella | 2899 |
| Hieracium auricula | 2857 |
| Hieracium lactucella | 2899 |
| Pilosella lactucella | 12147 |
| Hieracium murorum | 25659 |
| Hieracium murorum | 25659 |
| Hieracium sylvaticum | 2942 |
| Hieracium pilosella | 2923 |
| Hieracium pilosella | 2923 |
| Pilosella officinarum | 14992 |
| Hieracium piloselloides | 2924 |
| Hieracium piloselloides | 2924 |
| Pilosella piloselloides | 12226 |
| Hieracium rothianum | 25654 |
| Hieracium setigerum | 12219 |
| Hieracium subramosum | 10562 |
| depr. Hieracium subramosum | 94848 |
| Hippocrepis emerus | 21379 |
| Coronilla emerus | 1655 |
| Hornungia petraea | 2993 |
| Hornungia petraea | 2993 |
| Hutchinsia petraea | 22188 |
| Hypericum maculatum agg. | 3027 |
| Hypericum × desetangsii | 3029 |
| Hypericum maculatum | 20046 |
| Hypericum maculatum agg. | 3027 |
| Hypericum maculatum subsp. maculatum | 20047 |
| Hypericum perforatum | 3036 |
| Hypericum perforatum | 3036 |
| Hypericum perforatum subsp. angustifolium | 22208 |
| Hypericum tetrapterum | 3042 |
| Hypericum quadrangulum | 22214 |
| Hypericum tetrapterum | 3042 |
| Inula conyzae | 22227 |
| Inula conyza | 3069 |
| Inula conyzae | 22227 |
| Jasione laevis | 3117 |
| Jasione perennis | 22240 |
| Juncus alpinus | 22243 |
| Juncus alpinoarticulatus | 91853 |
| Juncus alpinus | 22243 |
| Juncus articulatus | 3136 |
| Juncus articulatus | 3136 |
| Juncus supinus | 22278 |
| Juncus bufonius agg. | 3140 |
| Juncus bufonius | 3141 |
| Juncus bufonius agg. | 3140 |
| Juncus compressus agg. | 3147 |
| Juncus compressus | 3148 |
| Juncus gerardii | 3149 |
| Juncus trifidus | 26603 |
| Juncus monanthos | 3165 |
| Juncus trifidus subsp. monanthos | 22281 |
| Juniperus communis | 3168 |
| Juniperus communis | 3168 |
| Juniperus communis subsp. communis | 3170 |
| Kobresia myosuroides | 22299 |
| Elyna myosuroides | 2102 |
| Kobresia myosuroides | 22299 |
| Koeleria pyramidata agg. | 3233 |
| Koeleria | 61398 |
| Koeleria cristata | 22305 |
| Koeleria gracilis | 22312 |
| Koeleria macrantha | 3235 |
| Koeleria pyramidata | 3236 |
| Koeleria pyramidata agg. | 3233 |
| Lamium galeobdolon agg. | 22337 |
| Galeobdolon argentatum | 21942 |
| Galeobdolon luteum | 20026 |
| Galeobdolon luteum agg. | 91498 |
| Galeobdolon montanum | 20027 |
| Lamiastrum galeobdolon | 3256 |
| Lamiastrum montanum | 3257 |
| Lamium argentatum | 24905 |
| Lamium galeobdolon | 22338 |
| Lamium galeobdolon agg. | 22337 |
| Lamium montanum | 22340 |
| Larix decidua | 3272 |
| Larix decidua | 3272 |
| Larix europaea | 10606 |
| Lathyrus linifolius | 3301 |
| Lathyrus linifolius | 3301 |
| Lathyrus montanus | 22390 |
| Ledum palustre | 3328 |
| Ledum palustre | 3328 |
| Rhododendron tomentosum | 28144 |
| Leontodon helveticus | 3346 |
| Leontodon helveticus | 3346 |
| Leontodon pyrenaicus subsp. helveticus | 7365 |
| Leontodon hispidus | 3347 |
| Leontodon hastilis | 25306 |
| Leontodon hispidus | 3347 |
| Leontodon hispidus subsp. hastilis | 22413 |
| Leontodon hispidus subsp. hispidus | 6837 |
| Leontodon saxatilis | 3354 |
| Leontodon saxatilis | 3354 |
| Leontodon taraxacoides | 22424 |
| Leucanthemum atratum agg. | 3375 |
| Leucanthemum atratum agg. | 3375 |
| Leucanthemum halleri | 3378 |
| Leucanthemum vulgare agg. | 3381 |
| Chrysanthemum leucanthemum | 21299 |
| Leucanthemum ircutianum | 3384 |
| Leucanthemum maximum | 3386 |
| Leucanthemum vulgare | 3387 |
| Leucanthemum vulgare agg. | 3381 |
| Linaria vulgaris agg. | 3429 |
| Linaria vulgaris | 3433 |
| Linaria vulgaris agg. | 3429 |
| Lindernia dubia | 3434 |
| Gratiola anagallidea | 13832 |
| Linum perenne agg. | 3454 |
| Linum austriacum | 3455 |
| Linum leonii | 3458 |
| Linum perenne | 26993 |
| Lithospermum arvense | 3475 |
| Buglossoides arvensis subsp. sibthorpiana | 20947 |
| Lithospermum arvense | 3475 |
| Lotus corniculatus agg. | 3505 |
| Lotus corniculatus | 3508 |
| Lotus corniculatus agg. | 3505 |
| Lotus corniculatus subsp. corniculatus | 22509 |
| Lotus corniculatus var. ciliatus | 25841 |
| Lotus tenuis | 3510 |
| Lotus pedunculatus | 22517 |
| Lotus pedunculatus | 22517 |
| Lotus uliginosus | 3516 |
| Luzula campestris agg. | 3530 |
| Luzula campestris | 3531 |
| Luzula campestris agg. | 3530 |
| Luzula campestris subsp. multiflora | 22525 |
| Luzula multiflora | 11817 |
| Luzula pallescens | 25388 |
| Luzula sudetica | 3536 |
| Luzula luzuloides | 3541 |
| Luzula albida | 3526 |
| Luzula luzuloides | 3541 |
| Luzula luzuloides subsp. rubella | 14424 |
| Luzula sylvatica | 3547 |
| Luzula sylvatica | 3547 |
| Luzula sylvatica subsp. sylvatica | 3549 |
| *Lycopersicon esculentum* | 3558 |
| *Lycopersicon esculentum* | 3558 |
| *Solanum lycopersicum* | 24088 |
| *Lycopus europaeus* | 3562 |
| *Lycopus europaeus* | 3562 |
| *Lycopus europaeus* subsp. *europaeus* | 3563 |
| *Malus sylvestris* agg. | 3580 |
| *Malus sylvestris* | 3582 |
| *Malus sylvestris* agg. | 3580 |
| *Matricaria* | 60765 |
| *Chamomilla* | 67025 |
| *Matricaria* | 60765 |
| *Matricaria discoidea* | 3597 |
| *Chamomilla suaveolens* | 21249 |
| *Matricaria discoidea* | 3597 |
| *Matricaria recutita* | 21248 |
| *Chamomilla recutita* | 21247 |
| *Matricaria recutita* | 21248 |
| *Medicago sativa* agg. | 3616 |
| *Medicago* × *varia* | 3620 |
| *Medicago falcata* | 3617 |
| *Medicago sativa* | 11820 |
| *Medicago sativa* agg. | 3616 |
| *Melampyrum pratense* | 3638 |
| *Melampyrum pratense* | 3638 |
| *Melampyrum pratense* subsp. *pratense* | 13853 |
| *Melica ciliata* agg. | 3643 |
| *Melica ciliata* | 3644 |
| *Melica ciliata* var. *nebrodensis* | 13015 |
| *Melica nutans* agg. | 3648 |
| *Melica nutans* | 3650 |
| *Melica nutans* agg. | 3648 |
| *Melica picta* | 3651 |
| *Mentha spicata* agg. | 3676 |
| *Mentha longifolia* | 3677 |
| *Mentha suaveolens* | 3680 |
| *Mercurialis perennis* agg. | 3691 |
| *Mercurialis ovata* | 3692 |
| *Mercurialis perennis* | 3694 |
| *Minuartia hybrida* | 3720 |
| *Alsine tenuifolia* | 25807 |
| *Minuartia rubra* | 22674 |
| *Minuartia fastigiata* | 3715 |
| *Minuartia rubra* | 22674 |
| *Minuartia verna* | 26607 |
| *Minuartia verna* | 26607 |
| *Minuartia verna* subsp. *gerardii* | 22675 |
| *Minuartia verna* subsp. *hercynica* | 6499 |
| *Molinia caerulea* agg. | 3756 |
| *Molinia* | 61408 |
| *Molinia arundinacea* | 3757 |
| *Molinia caerulea* | 3758 |
| *Monotropa hypopitys* agg. | 3762 |
| *Monotropa hypophegea* | 3763 |
| *Monotropa hypopitys* | 3764 |
| *Monotropa hypopitys* agg. | 3762 |
| *Montia fontana* | 3765 |
| *Montia fontana* | 3765 |
| *Montia fontana* subsp. *chondrosperma* | 3767 |
| *Mycelis muralis* | 3781 |
| *Lactuca muralis* | 22329 |
| *Mycelis muralis* | 3781 |
| *Myosotis arvensis* | 3782 |
| Myosotis arvensis | 3782 |
| Myosotis arvensis subsp. arvensis | 3783 |
| Myosotis discolor | 3785 |
| Myosotis discolor | 3785 |
| Myosotis versicolor | 22757 |
| Myosotis ramosissima | 3792 |
| Myosotis hispida | 22735 |
| Myosotis ramosissima | 3792 |
| Myosotis scorpioides agg. | 20051 |
| Myosotis caespitosa | 22731 |
| Myosotis laxa | 6500 |
| Myosotis laxiflora | 3788 |
| Myosotis multiflora | 12098 |
| Myosotis nemorosa | 3789 |
| Myosotis nemorosa subsp. brevisetacea | 22741 |
| Myosotis palustris | 22743 |
| Myosotis palustris agg. | 3786 |
| Myosotis palustris s. l. | 15315 |
| Myosotis scorpioides | 3791 |
| Myosotis scorpioides agg. | 20051 |
| Myosotis scorpioides subsp. scorpioides | 12104 |
| Myosotis stricta | 3804 |
| Myosotis micrantha | 22740 |
| Myosotis stricta | 3804 |
| Myosotis sylvatica agg. | 3793 |
| Myosotis alpestris | 3794 |
| Myosotis decumbens | 3795 |
| Myosotis sylvatica | 3802 |
| Nigritella nigra agg. | 3851 |
| Nigritella nigra | 7167 |
| Nigritella nigra agg. | 3851 |
| Odontites luteus | 3863 |
| Euphrasia lutea | 21768 |
| Odontites luteus | 3863 |
| Orthanthella lutea | 3863 |
| Odontites vernus agg. | 12023 |
| Odontites ruber | 13749 |
| Odontites vernus | 3866 |
| Odontites vernus agg. | 12023 |
| Odontites vulgaris | 3867 |
| Onobrychis viciifolia agg. | 3906 |
| Onobrychis arenaria | 3907 |
| Onobrychis sativa | 93791 |
| Onobrychis viciifolia | 3912 |
| Ononis spinosa agg. | 3920 |
| Ononis procurrens | 22831 |
| Ononis repens | 3922 |
| Ononis repens subsp. procurrens | 3923 |
| Ononis spinosa | 3925 |
| Ononis spinosa agg. | 3920 |
| Ononis spinosa subsp. maritima | 22837 |
| Ophrys holoserica | 3955 |
| Ophrys fuciflora | 22852 |
| Ophrys holoserica | 3955 |
| Ophrys insectifera | 3956 |
| Ophrys insectifera | 3956 |
| Ophrys muscifera | 22859 |
| Oreopteris limbosperma | 22909 |
| Dryopteris montana | 21597 |
| Lastrea limbosperma | 22377 |
| Oreopteris limbosperma | 22909 |
| Thelypteris limbosperma | 5910 |
| Orthilia secunda | 4054 |
| Orthilia secunda | 4054 |
| Pyrola secunda | 23440 |
| Oxalis stricta | 22973 |
| Oxalis fontana | 4065 |
| Oxalis stricta | 22973 |
| Oxytropis montana agg. | 4080 |
| Oxytropis jacquinii | 4084 |
| Oxytropis montana agg. | 4080 |
| Papaver dubium | 10700 |
| Papaver dubium | 10700 |
| Papaver dubium subsp. lecoqii | 22999 |
| Pedicularis palustris | 4158 |
| Pedicularis palustris | 4158 |
| Pedicularis palustris subsp. palustris | 13828 |
| Persicaria amphibia | 23027 |
| Persicaria amphibia | 23027 |
| Polygonum amphibium | 4413 |
| Persicaria dubia | 11734 |
| Persicaria dubia | 11734 |
| Persicaria mitis | 23041 |
| Polygonum mite | 4429 |
| Persicaria hydropiper | 23032 |
| Persicaria hydropiper | 23032 |
| Polygonum hydropiper | 4422 |
| Persicaria lapathifolia | 23034 |
| Persicaria lapathifolia | 23034 |
| Persicaria lapathifolia subsp. pallida | 13950 |
| Polygonum lapathifolium | 4423 |
| Persicaria maculosa | 23038 |
| Persicaria maculosa | 23038 |
| Polygonum persicaria | 4435 |
| Persicaria minor | 23039 |
| Persicaria minor | 23039 |
| Polygonum minus | 4428 |
| Petrorhagia prolifera agg. | 4185 |
| Petrorhagia prolifera | 4187 |
| Tunica prolifera | 24402 |
| Petrorhagia saxifraga | 4189 |
| Petrorhagia saxifraga | 4189 |
| Tunica saxifraga | 24403 |
| Phegopteris connectilis | 23065 |
| Gymnocarpium phegopteris | 12299 |
| Phegopteris connectilis | 23065 |
| Thelypteris phegopteris | 5912 |
| Phleum alpinum agg. | 4215 |
| Phleum alpinum | 25110 |
| Phleum alpinum agg. | 4215 |
| Phleum phleoides | 4222 |
| Phleum boehmeri | 23073 |
| Phleum phleoides | 4222 |
| Phleum pratense agg. | 4223 |
| Phleum bertolonii | 4224 |
| Phleum pratense | 4225 |
| Phleum pratense agg. | 4223 |
| Phleum pratense subsp. pratense | 27013 |
| Phragmites australis | 4229 |
| Phragmites australis | 4229 |
| Phragmites communis | 23084 |
| Phyteuma orbiculare | 26610 |
| Phyteuma orbiculare | 26610 |
| Phyteuma orbiculare subsp. orbiculare | 24940 |
| Phyteuma spicatum | 4264 |
| Phyteuma spicatum | 4264 |
| Phyteuma spicatum subsp. occidentale | 14758 |
| Phyteuma spicatum subsp. spicatum | 4266 |
| Phyteuma spicatum var. coeruleum | 23091 |
| Picea abies | 4269 |
| Picea abies | 4269 |
| Picea excelsa | 23106 |
| Picris hieracioides | 4274 |
| Picris hieracioides | 4274 |
| Picris hieracioides agg. | 4272 |
| Pimpinella major | 4277 |
| Pimpinella major | 4277 |
| Pimpinella major subsp. major | 23108 |
| Pimpinella saxifraga agg. | 4279 |
| Pimpinella saxifraga | 4282 |
| Pimpinella saxifraga agg. | 4279 |
| Plantago lanceolata | 4320 |
| Plantago lanceolata | 4320 |
| Plantago lanceolata subsp. sphaerostachya | 23153 |
| Plantago lanceolata var. sphaerostachya | 25683 |
| Plantago major | 4321 |
| Plantago intermedia | 23150 |
| Plantago major | 4321 |
| Plantago major subsp. intermedia | 4322 |
| Plantago major subsp. major | 4323 |
| Plantago maritima agg. | 4325 |
| Plantago alpina | 4326 |
| Plantago maritima | 27766 |
| Plantago strictissima | 23161 |
| Plantago media agg. | 4332 |
| Plantago media | 4333 |
| Plantago media agg. | 4332 |
| Poa annua agg. | 4343 |
| Poa annua | 4344 |
| Poa annua agg. | 4343 |
| Poa supina | 4345 |
| Poa pratensis agg. | 4366 |
| Poa angustifolia | 4367 |
| Poa humilis | 7372 |
| Poa pratensis | 4368 |
| Poa pratensis agg. | 4366 |
| Poa pratensis subsp. angustifolia | 23193 |
| Poa trivialis | 26611 |
| Poa trivialis | 26611 |
| Poa trivialis agg. | 4372 |
| Poa trivialis subsp. trivialis | 7161 |
| Polygala amara agg. | 4390 |
| Polygala amara | 4391 |
| Polygala amara agg. | 4390 |
| Polygala amara subsp. amarella | 23205 |
| Polygala amarella | 4394 |
| Polygala vulgaris | 4405 |
| Polygala oxyptera | 23212 |
| Polygala vulgaris | 4405 |
| Polygala vulgaris s. str. | 10745 |
| Polygala vulgaris subsp. oxyptera | 4406 |
| Polygala vulgaris subsp. vulgaris | 4407 |
| Polygonatum odoratum | 4410 |
| Polygonatum odoratum | 4410 |
| Polygonatum officinale | 27134 |
| Polygonum aviculare agg. | 4415 |
| Polygonum aviculare | 4417 |
| Polygonum aviculare agg. | 4415 |
| Populus x canadensis | 50080 |
| Populus × canadensis | 50080 |
| Populus × euamericana | 10760 |
| Potentilla argentea agg. | 4490 |
| Potentilla argentea | 4491 |
| Potentilla argentea agg. | 4490 |
| Potentilla erecta | 4511 |
| Potentilla erecta | 4511 |
| Potentilla tormentilla | 23354 |
| Potentilla heptaphylla agg. | 4515 |
| Potentilla heptaphylla | 4517 |
| Potentilla rubens | 23350 |
| Potentilla incana | 25757 |
| Potentilla arenaria | 4539 |
| Potentilla cinerea | 25853 |
| Potentilla cinerea subsp. incana | 15377 |
| Potentilla incana | 25757 |
| Potentilla × subarenaria | 50084 |
| Potentilla palustris | 4528 |
| Comarum palustre | 21352 |
| Potentilla palustris | 4528 |
| Potentilla verna agg. | 4538 |
| Potentilla neumanniana | 4541 |
| Potentilla pusilla | 4542 |
| Potentilla tabernaemontani | 20053 |
| Potentilla verna | 23355 |
| Potentilla verna agg. | 4538 |
| Potentilla verna subsp. vulgaris | 28077 |
| Primula veris | 4570 |
| Primula officinalis | 23378 |
| Primula veris | 4570 |
| Primula veris subsp. veris | 4575 |
| Primula × digenea | 92537 |
| Primula elatior × vulgaris | 92534 |
| Primula × media | 50085 |
| Primula elatior × veris | 92533 |
| Pritzelago alpina | 26680 |
| Hutchinsia alpina | 3000 |
| Pritzelago alpina | 26680 |
| Prunus avium | 4582 |
| Cerasus avium | 21216 |
| Prunus avium | 4582 |
| Prunus avium subsp. avium | 24942 |
| Prunus cerasus agg. | 4585 |
| Cerasus vulgaris | 21220 |
| Prunus cerasus | 4586 |
| Prunus mahaleb | 4592 |
| Cerasus mahaleb | 21219 |
| Prunus mahaleb | 4592 |
| Prunus padus | 4593 |
| Padus avium | 22980 |
| Prunus padus | 4593 |
| Prunus spinosa agg. | 4597 |
| Prunus × fruticans | 4598 |
| Prunus spinosa | 4599 |
| Prunus spinosa subsp. fruticans | 23396 |
| Pseudolysimachion longifolium | 23400 |
| Pseudolysimachion longifolium | 23400 |
| Veronica longifolia | 6261 |
| Pseudolysimachion spicatum | 23404 |
| Pseudolysimachion spicatum | 23404 |
| Veronica spicata | 6286 |
| Pseudorchis albida | 4601 |
| Leucorchis albida | 3390 |
| Pseudorchis albida | 4601 |
| Pulmonaria officinalis agg. | 4636 |
| Pulmonaria obscura | 4637 |
| Pulmonaria officinalis | 4638 |
| Pulmonaria officinalis agg. | 4636 |
| Pulsatilla alpina | 15385 |
| Pulsatilla alpina agg. | 4642 |
| Pulsatilla vulgaris | 26617 |
| Anemone pulsatilla | 20428 |
| Anemone pulsatilla subsp. grandis | 20429 |
| Pulsatilla vulgaris | 26617 |
| Pulsatilla vulgaris agg. | 4656 |
| Pulsatilla vulgaris subsp. grandis | 23437 |
| Pyrus communis agg. | 4669 |
| Pyrus communis | 6936 |
| Pyrus communis agg. | 4669 |
| Pyrus communis subsp. pyraeaster | 23449 |
| Pyrus pyraeaster | 4671 |
| Quercus petraea agg. | 4678 |
| Quercus petraea | 4680 |
| Quercus sessiliflora | 23468 |
| Quercus robur | 4685 |
| Quercus pedunculata | 23466 |
| Quercus robur | 4685 |
| Ranunculus acris agg. | 4689 |
| Ranunculus acris | 4690 |
| Ranunculus acris agg. | 4689 |
| Ranunculus aquatilis agg. | 4697 |
| Ranunculus aquatilis | 4698 |
| Ranunculus aquatilis agg. | 4697 |
| Ranunculus peltatus | 12030 |
| Ranunculus auricomus agg. | 4709 |
| Ranunculus auricomus | 11978 |
| Ranunculus auricomus agg. | 4709 |
| Ranunculus auricomus s. l. | 4710 |
| Ranunculus biflorus | 27241 |
| Ranunculus kochii | 10821 |
| Ranunculus rectus | 6956 |
| Ranunculus bulbosus | 4717 |
| Ranunculus bulbosus | 4717 |
| Ranunculus bulbosus subsp. bulbosus | 12015 |
| Ranunculus ficaria | 4721 |
| Ficaria verna | 21893 |
| Ranunculus ficaria | 4721 |
| Ranunculus ficaria subsp. bulbilifer | 4722 |
| Ranunculus flammula agg. | 4726 |
| Ranunculus flammula | 4727 |
| Ranunculus flammula agg. | 4726 |
| Ranunculus reptans | 4728 |
| Ranunculus montanus agg. | 4743 |
| Ranunculus breynius | 23477 |
| Ranunculus montanus | 4747 |
| Ranunculus montanus agg. | 4743 |
| Ranunculus polyanthemus agg. | 4758 |
| Ranunculus nemorosus | 4759 |
| Ranunculus nemorosus subsp. polyanthemophyllus | 23501 |
| Ranunculus polyanthemoides | 4760 |
| Ranunculus polyanthemophyllus | 4761 |
| Ranunculus polyanthemus | 4762 |
| Ranunculus polyanthemus agg. | 4758 |
| Raphanus raphanistrum agg. | 4778 |
| Raphanus raphanistrum | 4780 |
| Raphanus sativus | 4781 |
| Rhinanthus angustifolius | 23544 |
| Rhinanthus angustifolius | 23544 |
| Rhinanthus angustifolius subsp. angustifolius | 23545 |
| Rhinanthus angustifolius subsp. grandiflorus | 6569 |
| Rhinanthus serotinus | 4832 |
| Rhinanthus aristatus agg. | 4817 |
| Rhinanthus aristatus | 23551 |
| Rhinanthus glacialis | 4819 |
| Rhinanthus minor | 4828 |
| Alectorolophus minor | 20304 |
| Rhinanthus minor | 4828 |
| Ribes rubrum agg. | 4850 |
| Ribes rubrum | 4851 |
| Ribes rubrum agg. | 4850 |
| Rosa canina agg. s. l. | 94740 |
| Rosa canina | 26665 |
| Rosa canina agg. | 4872 |
| Rosa canina agg. s. l. | 94740 |
| Rosa corymbifera | 26666 |
| Rosa obtusifolia | 4897 |
| Rosa rubiginosa agg. s. l. | 92752 |
| Rosa elliptica | 4887 |
| Rosa micrantha | 4893 |
| Rosa rubiginosa | 4902 |
| Rosa rubiginosa agg. s. l. | 92752 |
| Rosa spinosissima | 23649 |
| Rosa pimpinellifolia | 4899 |
| Rosa tomentosa agg. s. l. | 92754 |
| Rosa tomentosa | 4912 |
| Rosa tomentosa agg. | 4907 |
| Rosa villosa agg. | 4908 |
| Rubus canescens | 4952 |
| Rubus canescens | 4952 |
| Rubus tomentosus | 25709 |
| Rubus grabowskii | 7078 |
| Rubus thyrsanthus | 5056 |
| Rubus latiarcuatus | 24749 |
| Rubus vulgaris var. mollis | 11374 |
| Rubus montanus | 24778 |
| Rubus candicans | 4951 |
| Rubus montanus | 24778 |
| Rubus plicatus | 5024 |
| Rubus affinis | 4931 |
| Rubus plicatus | 5024 |
| Rubus sect. Rubus | 11351 |
| Rubus fruticosus s. l. | 11886 |
| Rubus sect. Rubus | 11351 |
| Rumex acetosa agg. | 92799 |
| Rumex acetosa | 5073 |
| Rumex thyrsiflorus | 5108 |
| Rumex acetosella | 26619 |
| Rumex acetosella | 26619 |
| Rumex acetosella agg. | 5074 |
| Rumex acetosella subsp. acetosella | 26618 |
| Rumex acetosella subsp. tenuifolius | 27201 |
| Rumex tenuifolius | 5077 |
| Rumex arifolius | 20080 |
| Rumex alpestris | 5078 |
| Rumex arifolius | 20080 |
| Rumex pseudoalpinus | 23661 |
| Rumex alpinus | 5079 |
| Rumex × pratensis | 50156 |
| Rumex × pratensis | 50156 |
| Rumex crispus × obtusifolius | 92814 |
| Salix fragilis agg. | 5167 |
| Salix × rubens | 5169 |
| Salix fragilis | 5168 |
| Salix myrsinifolia | 5185 |
| Salix myrsinifolia | 5185 |
| Salix nigricans | 23730 |
| Salix repens agg. | 27193 |
| Salix repens | 27194 |
| Salix repens agg. | 27193 |
| Salix repens s. l. | 5191 |
| Salix repens subsp. repens | 5193 |
| Salix retusa agg. | 5196 |
| Salix retusa | 5198 |
| Salix retusa agg. | 5196 |
| Salix serpillifolia | 5199 |
| Salix × multinervis | 27195 |
| Salix × multinervis | 27195 |
| Salix aurita × cinerea | 92857 |
| Salix × smithiana | 27196 |
| Salix × smithiana | 27196 |
| Salix caprea × viminalis | 92873 |
| Salix × wimmeriana | 50108 |
| Salix caprea × purpurea | 92872 |
| Salsola kali | 5208 |
| Salsola kali subsp. tragus | 5211 |
| Salsola tragus | 23749 |
| Sanguisorba minor | 5231 |
| Sanguisorba minor | 5231 |
| Sanguisorba minor subsp. minor | 5232 |
| Scabiosa columbaria agg. | 5337 |
| Scabiosa columbaria | 5338 |
| Scabiosa columbaria agg. | 5337 |
| Scabiosa lucida | 5342 |
| Scleranthus annuus agg. | 5376 |
| Scleranthus annuus | 5377 |
| Scleranthus annuus agg. | 5376 |
| Scleranthus polycarpus | 5378 |
| Scleranthus × intermedius | 92980 |
| Scleranthus annuus × perennis | 92978 |
| Securigera varia | 21382 |
| Coronilla varia | 1662 |
| Securigera varia | 21382 |
| Sedum rupestre agg. | 5429 |
| Sedum reflexum | 5433 |
| Sedum rupestre | 23907 |
| Sedum rupestre subsp. reflexum | 23910 |
| Sedum sexangulare | 5437 |
| Sedum boloniense | 23894 |
| Sedum mite | 23901 |
| Sedum sexangulare | 5437 |
| Sedum telephium agg. | 5440 |
| Hylotelephium maximum | 7126 |
| Sedum maximum | 5441 |
| Sedum telephium | 27746 |
| Sedum telephium agg. | 5440 |
| Sedum telephium subsp. maximum | 23912 |
| Senecio alpinus | 5467 |
| Senecio alpinus | 5467 |
| Senecio cordatus | 23945 |
| Senecio aquaticus agg. | 5468 |
| Senecio aquaticus | 5469 |
| Senecio aquaticus agg. | 5468 |
| *Senecio erraticus* | 5470 |
| *Senecio incanus* | 5485 |
| *Senecio carniolicus* | 23942 |
| *Senecio jacobaea* | 5494 |
| *Jacobaea vulgaris* | 25789 |
| *Senecio jacobaea* | 5494 |
| *Senecio nemorensis* agg. | 5496 |
| *Senecio fuchsii* | 5498 |
| *Senecio hercynicus* | 5499 |
| *Senecio nemorensis* | 94860 |
| *Senecio nemorensis* agg. | 5496 |
| *Senecio ovatus* | 23947 |
| *Senecio ovatus* subsp. *alpestris* | 7006 |
| *Serratula tinctoria* | 27047 |
| *Serratula tinctoria* | 27047 |
| *Serratula tinctoria* agg. | 5524 |
| *Seseli libanotis* | 5540 |
| *Libanotis montana* | 22460 |
| *Libanotis sibirica* | 22463 |
| *Seseli libanotis* | 5540 |
| *Sesleria varia* agg. | 5558 |
| *Sesleria albicans* | 23996 |
| *Sesleria albicans* subsp. *albicans* | 27051 |
| *Sesleria albicans* var. *albicans* | 15414 |
| *Sesleria caerulea* | 25028 |
| *Sesleria caerulea* subsp. *calcaria* | 13441 |
| *Sesleria varia* | 24991 |
| *Setaria pumila* | 24008 |
| *Setaria glauca* | 5563 |
| *Setaria pumila* | 24008 |
| *Silene* | 60691 |
| *Lychnis* | 60681 |
| *Silene* | 60691 |
| *Silene acaulis* agg. | 5578 |
| *Silene acaulis* | 5579 |
| *Silene acaulis* agg. | 5578 |
| *Silene coronaria* | 11625 |
| *Lychnis coronaria* | 3551 |
| *Silene flos-cuculi* | 24031 |
| *Lychnis flos-cuculi* | 3552 |
| *Silene flos-cuculi* | 24031 |
| *Silene latifolia* | 7012 |
| *Melandrium album* | 22627 |
| *Silene alba* | 5581 |
| *Silene latifolia* | 7012 |
| *Silene latifolia* subsp. *alba* | 22546 |
| *Silene pratensis* | 20058 |
| *Silene noctiflora* | 5603 |
| *Melandrium noctiflorum* | 22630 |
| *Silene noctiflora* | 5603 |
| *Silene nutans* agg. | 5604 |
| *Silene nutans* | 5606 |
| *Silene nutans* agg. | 5604 |
| *Silene viscaria* | 11627 |
| *Lychnis viscaria* | 3554 |
| *Silene viscaria* | 11627 |
| *Viscaria vulgaris* | 24569 |
| *Silene vulgaris* | 5629 |
| *Silene cucubalus* | 24029 |
| *Silene inflata* | 25277 |
| *Silene vulgaris* | 5629 |
| *Silene vulgaris* agg. | 20067 |
| *Silene vulgaris* subsp. *vulgaris* | 5634 |
| Silene vulgaris var. humilis | 24046 |
| Soldanella alpicola | 24100 |
| Soldanella pusilla s. l. | 5678 |
| Solidago virgaurea | 5682 |
| Solidago virgaurea | 5682 |
| Solidago virgaurea subsp. virgaurea | 5684 |
| Sonchus asper | 5690 |
| Sonchus asper | 5690 |
| Sonchus asper subsp. asper | 5691 |
| Sorbus aria agg. | 5696 |
| Sorbus aria | 5697 |
| Sorbus aria agg. | 5696 |
| Sorbus aucuparia | 5700 |
| Sorbus aucuparia | 5700 |
| Sorbus aucuparia subsp. aucuparia | 5701 |
| Spergula pentandra agg. | 5730 |
| Spergula morisonii | 5731 |
| Spergula vernalis | 24154 |
| Spergularia media | 5734 |
| Spergularia maritima | 24145 |
| Spergularia media | 5734 |
| Stellaria alsine | 5769 |
| Stellaria alsine | 5769 |
| Stellaria uliginosa | 5782 |
| Stellaria aquatica | 24917 |
| Myosoton aquaticum | 3805 |
| Stellaria aquatica | 24917 |
| Stellaria media agg. | 5774 |
| Stellaria media | 5775 |
| Stellaria media agg. | 5774 |
| Stellaria pallida | 5777 |
| Stellaria nemorum | 5778 |
| Stellaria nemorum | 5778 |
| Stellaria nemorum subsp. nemorum | 5780 |
| Stellaria palustris | 11573 |
| Stellaria glauca | 24184 |
| Stellaria palustris | 11573 |
| Stipa calamagrostis | 24197 |
| Achnatherum calamagrostis | 47 |
| Stipa pennata agg. | 5787 |
| Stipa joannis | 5797 |
| Stipa pennata | 24203 |
| Stipa pennata var. mediterranea | 13411 |
| Stipa pulcherrima | 13746 |
| Symphytum officinale | 26629 |
| Symphytum officinale | 26629 |
| Symphytum officinale agg. | 5823 |
| Tanacetum corymbosum agg. | 5841 |
| Chrysanthemum corymbosum | 21294 |
| Tanacetum corymbosum | 5843 |
| Tanacetum parthenium | 5845 |
| Chrysanthemum parthenium | 21315 |
| Tanacetum parthenium | 5845 |
| Tanacetum vulgare | 5846 |
| Chrysanthemum vulgare | 21314 |
| Tanacetum vulgare | 5846 |
| Taraxacum sect. Alpina et Hamata et Ruderalia | 93219 |
| Taraxacum sect. Alpina | 7424 |
| Taraxacum officinale agg. | 7442 |
| Taraxacum sect. Alpina et Hamata et Ruderalia | 93219 |
| Taraxacum sect. Ruderalia | 7434 |
| Taraxacum officinale | 14513 |
| Taraxacum sect. Erythrosperma | 7428 |
| Taraxacum erythrospermum | 14460 |
| Taraxacum laevigatum | 14457 |
| Taraxacum levigatum agg. | 14458 |
| Taraxacum sect. Erythrosperma | 7428 |
| Taraxacum sect. Palustria | 7433 |
| Taraxacum palustre | 25514 |
| Taraxacum sect. Palustria | 7433 |
| Tephroseris helenitis | 23949 |
| Senecio helenitis | 5482 |
| Tephroseris helenitis | 23949 |
| Tephroseris integrifolia | 23953 |
| Senecio integrifolius | 5492 |
| Tephroseris integrifolia | 23953 |
| Teucrium scorodonia | 5889 |
| Teucrium scorodonia | 5889 |
| Teucrium scorodonia subsp. scorodonia | 5891 |
| Thalictrum minus agg. | 5896 |
| Thalictrum minus | 5898 |
| Thalictrum minus agg. | 5896 |
| Thalictrum minus subsp. saxatile | 5902 |
| Thlaspi caerulescens | 5928 |
| Noccaea caerulescens | 22785 |
| Thlaspi cepaeifolium | 25000 |
| Thlaspi rotundifolium | 5940 |
| Thlaspi perfoliatum | 5938 |
| Microthlaspi perfoliatum | 22667 |
| Thlaspi perfoliatum | 5938 |
| Thymus praecox agg. | 5955 |
| Thymus praecox | 5957 |
| Thymus praecox agg. | 5955 |
| Thymus praecox subsp. polytrichus | 5959 |
| Thymus praecox subsp. praecox | 5960 |
| Thymus serpyllum subsp. praecox | 13239 |
| Thymus pulegioides agg. | 5964 |
| Thymus pulegioides | 5965 |
| Thymus pulegioides agg. | 5964 |
| Thymus pulegioides subsp. pulegioides | 7053 |
| Thymus serpyllum subsp. chamaedrys | 13255 |
| Thymus serpyllum agg. | 6456 |
| Thymus angustifolius | 24265 |
| Thymus serpyllum | 5966 |
| Thymus serpyllum agg. | 6456 |
| Tragopogon pratensis | 7141 |
| Tragopogon minor | 5999 |
| Tragopogon orientalis | 6000 |
| Tragopogon pratensis | 7141 |
| Tragopogon pratensis agg. | 5998 |
| Tragopogon pratensis subsp. orientalis | 24325 |
| Tragopogon pratensis subsp. pratensis | 7100 |
| Trichophorum cespitosum | 27072 |
| Trichophorum cespitosum | 27072 |
| Trichophorum cespitosum subsp. germanicum | 24330 |
| Trifolium dubium agg. | 6028 |
| Trifolium dubium | 6029 |
| Trifolium dubium agg. | 6028 |
| Trifolium minus | 24352 |
| Trifolium pratense | 6057 |
| Trifolium pratense | 6057 |
| Trifolium pratense subsp. pratense | 6061 |
| Tripleurospermum maritimum agg. | 6096 |
| Matricaria inodora | 22000 |
| Matricaria maritima | 22580 |
| Matricaria perforata | 22003 |
| Tripleurospermum inodorum | 20061 |
| Tripleurospermum maritimum | 6098 |
| Tripleurospermum perforatum | 6097 |
| Ulmus glabra | 6137 |
| Ulmus glabra | 6137 |
| Ulmus montana | 24424 |
| Ulmus scabra | 24425 |
| Ulmus minor agg. | 6139 |
| Ulmus campestris | 24420 |
| Ulmus minor | 6140 |
| Utricularia minor agg. | 6152 |
| Utricularia minor | 6154 |
| Utricularia minor agg. | 6152 |
| Vaccinium oxycoccos agg. | 6161 |
| Oxycoccus palustris | 4070 |
| Vaccinium oxycoccos | 6163 |
| Vaccinium oxycoccos agg. | 6161 |
| Vaccinium uliginosum | 6166 |
| Vaccinium uliginosum | 6166 |
| Vaccinium uliginosum agg. | 6164 |
| Valeriana officinalis agg. | 6177 |
| Valeriana officinalis | 6178 |
| Valeriana officinalis agg. | 6177 |
| Valeriana pratensis | 6179 |
| Valeriana procurrens | 6180 |
| Valeriana wallrothii | 6183 |
| Veronica agrestis agg. | 93432 |
| Veronica agrestis | 6226 |
| Veronica persica | 6271 |
| Veronica polita | 6272 |
| Veronica anagallis-aquatica agg. | 6231 |
| Veronica anagallis-aquatica | 6232 |
| Veronica anagalloides | 6233 |
| Veronica catenata | 6234 |
| Veronica austriaca agg. | 6237 |
| Veronica austriaca | 13751 |
| Veronica austriaca subsp. dentata | 24471 |
| Veronica austriaca subsp. teucrium | 25019 |
| Veronica prostrata | 13869 |
| Veronica teucrium | 6243 |
| Veronica chamaedrys | 13752 |
| Veronica chamaedrys | 13752 |
| Veronica chamaedrys agg. | 6248 |
| Veronica hederifolia | 26766 |
| Veronica hederifolia | 26766 |
| Veronica hederifolia agg. | 6257 |
| Veronica hederifolia subsp. hederifolia | 7101 |
| Veronica sublobata | 6259 |
| Veronica verna agg. | 6292 |
| Veronica dillenii | 6293 |
| Veronica verna | 6294 |
| Veronica verna agg. | 6292 |
| Vicia cracca agg. | 6301 |
| Vicia cracca | 6302 |
| Vicia cracca agg. | 6301 |
| Vicia tenuifolia | 6306 |
| Vicia sativa agg. | 6328 |
| Vicia angustifolia | 6329 |
| Vicia sativa | 6334 |
| Vicia sativa agg. | 6328 |
| Vicia sativa subsp. nigra | 24521 |
| Vicia sativa var. nigra | 24518 |
| Vicia tetrasperma agg. | 6338 |
| Vicia tenuissima | 6340 |
| Vicia tetrasperma | 6341 |
| Vicia tetrasperma agg. | 6338 |
| Vincetoxicum hirundinaria agg. | 6349 |
| Cynanchum vincetoxicum | 21465 |
| Vincetoxicum hirundinaria | 6351 |
| Vincetoxicum officinale | 24536 |
| Viola canina agg. | 24994 |
| Viola canina | 6367 |
| Viola canina subsp. canina | 6368 |
| Viola persicifolia | 6386 |
| Viola persicifolia | 6386 |
| Viola stagnina | 6394 |
| Viola silvatica agg. | 94746 |
| Viola × bavarica | 7420 |
| Viola reichenbachiana | 6390 |
| Viola reichenbachiana × riviniana | 93487 |
| Viola riviniana | 6391 |
| Viola silvatica | 24559 |
| Viola tricolor agg. | 6397 |
| Viola arvensis | 6398 |
| Viola arvensis × tricolor | 93457 |
| Viola arvensis subsp. arvensis | 7208 |
| Viola tricolor | 6402 |
| Viola tricolor subsp. tricolor | 6405 |
| Viola tricolor var. arvensis | 12796 |
| x Festulolium krasanii | 93548 |
| Festuca arundinacea × Lolium multiflorum | 91392 |

**Table 6:**
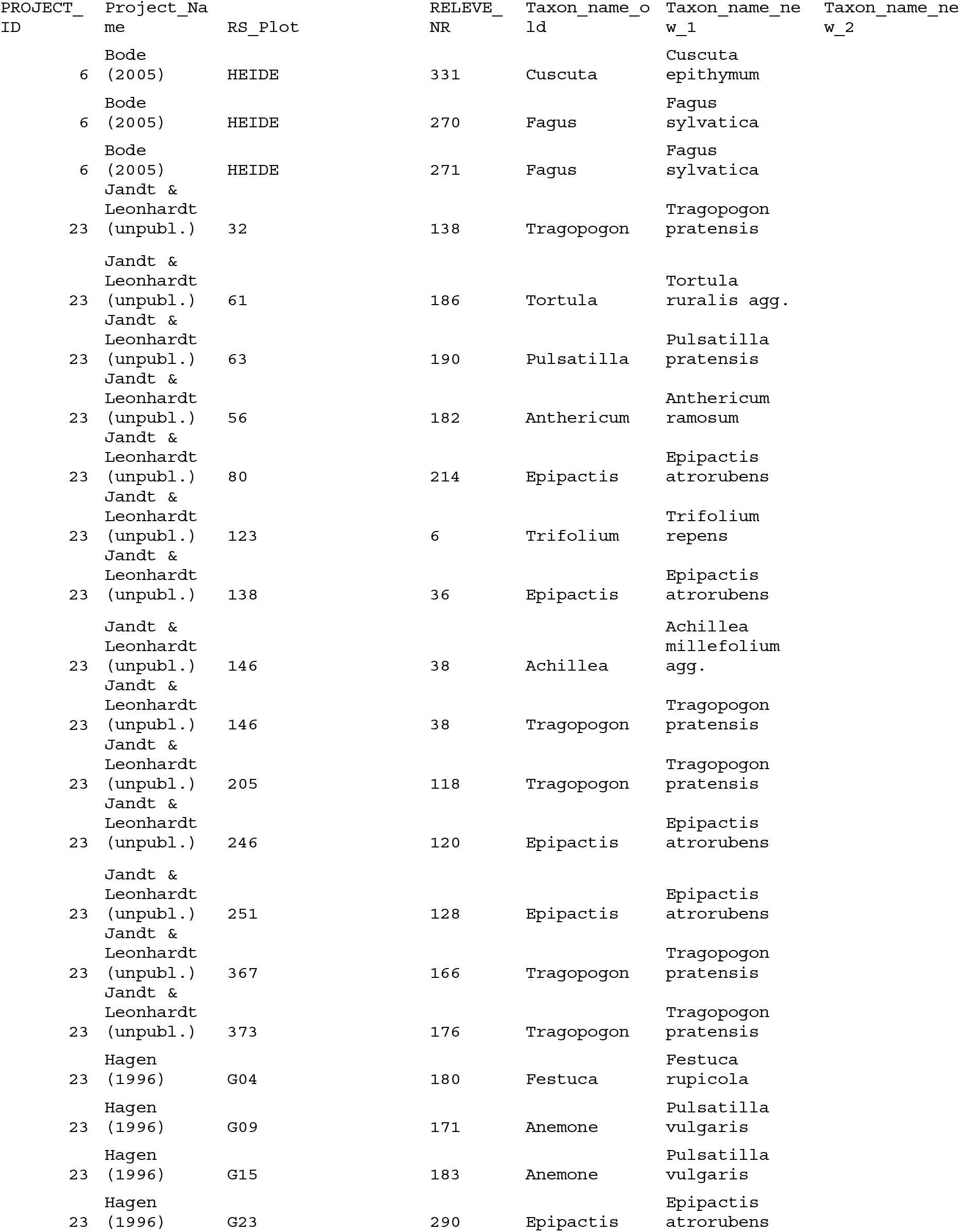
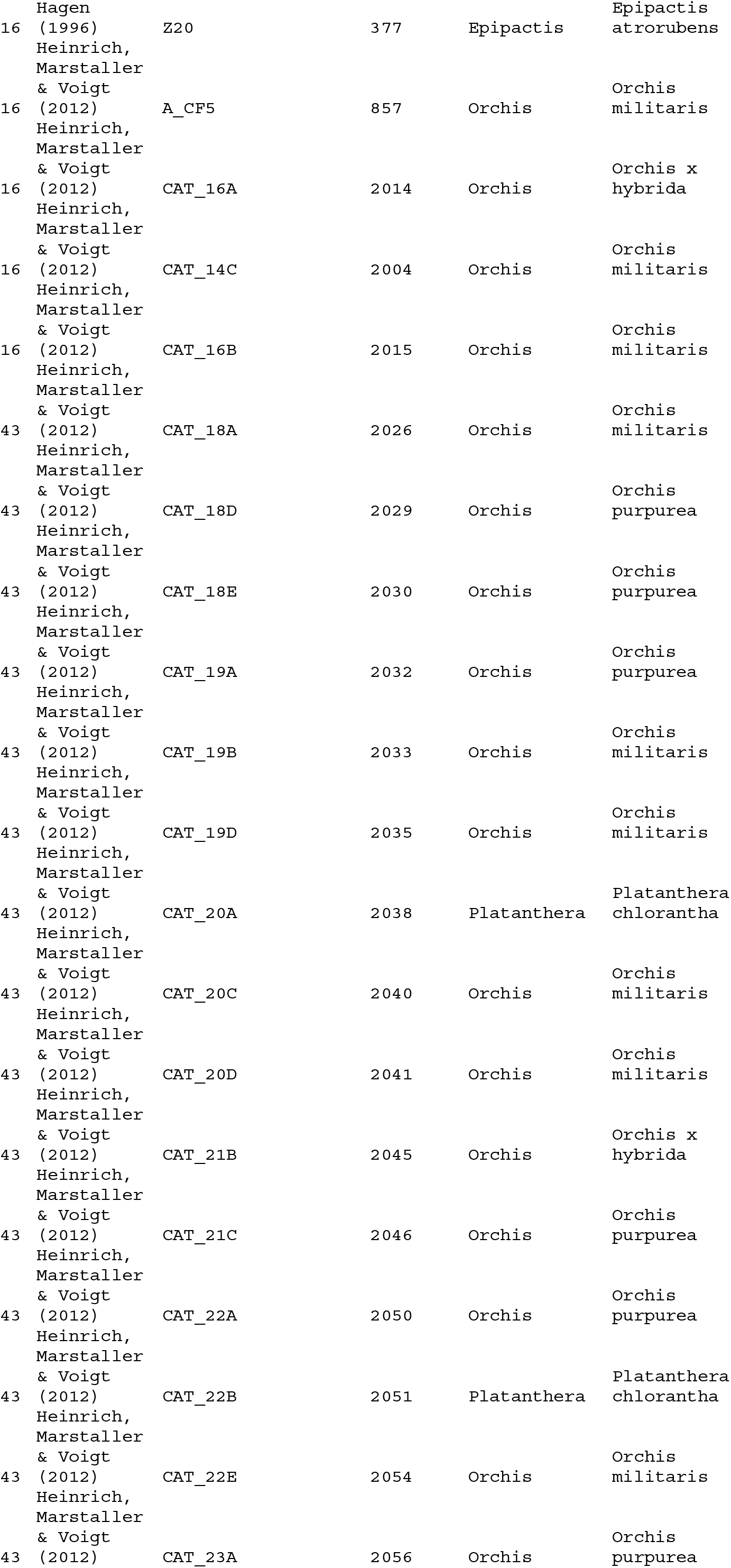
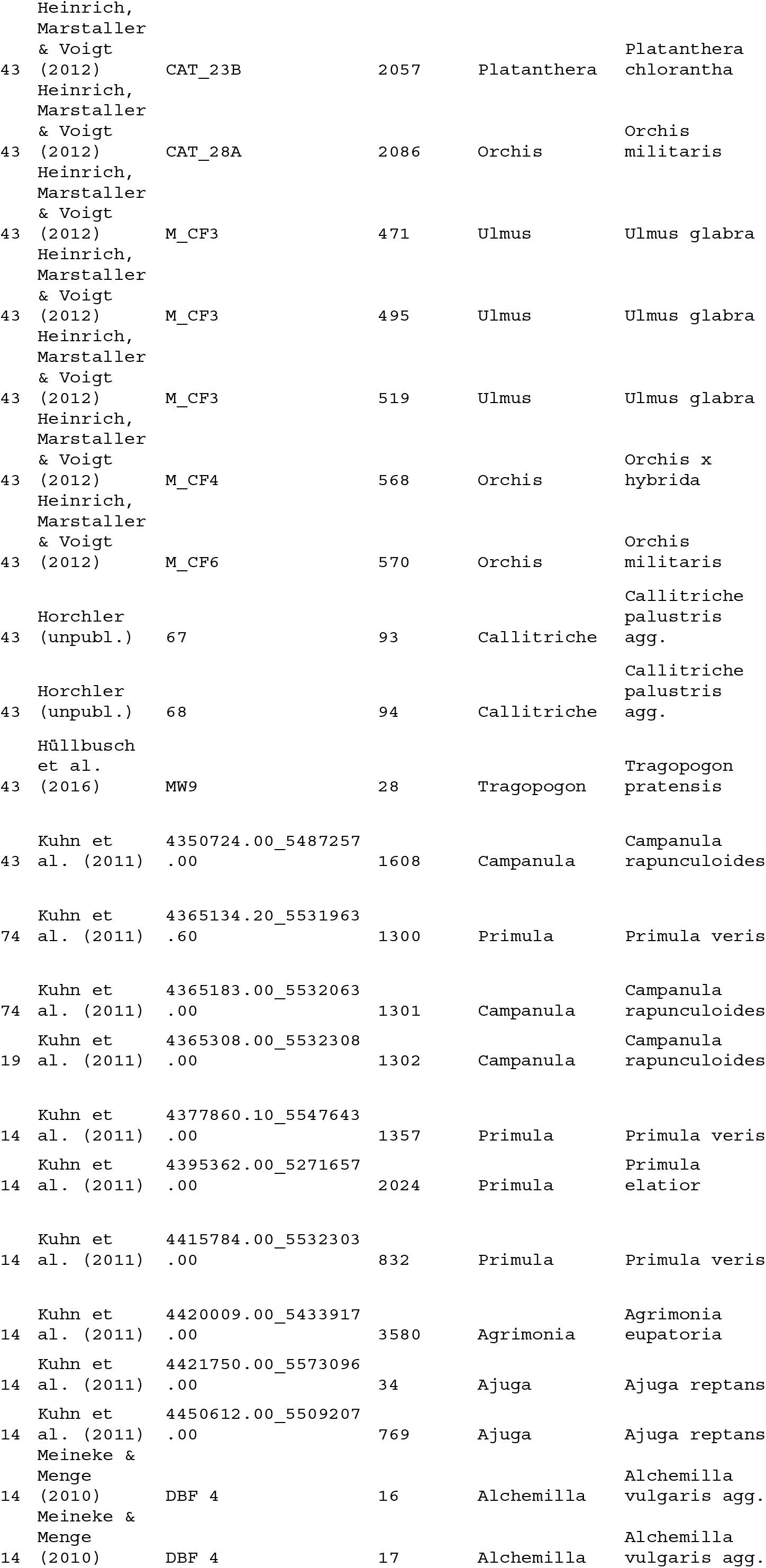
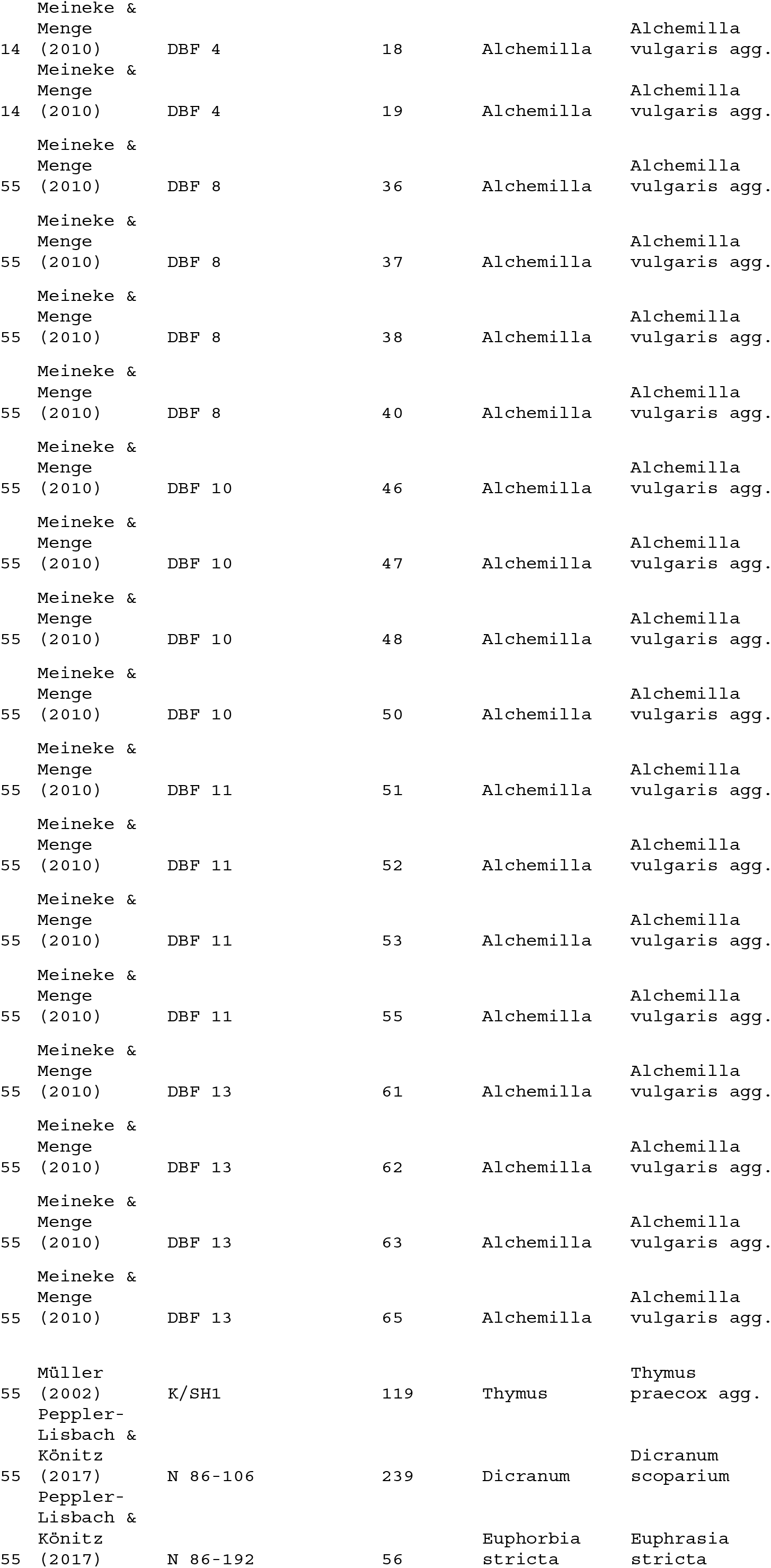
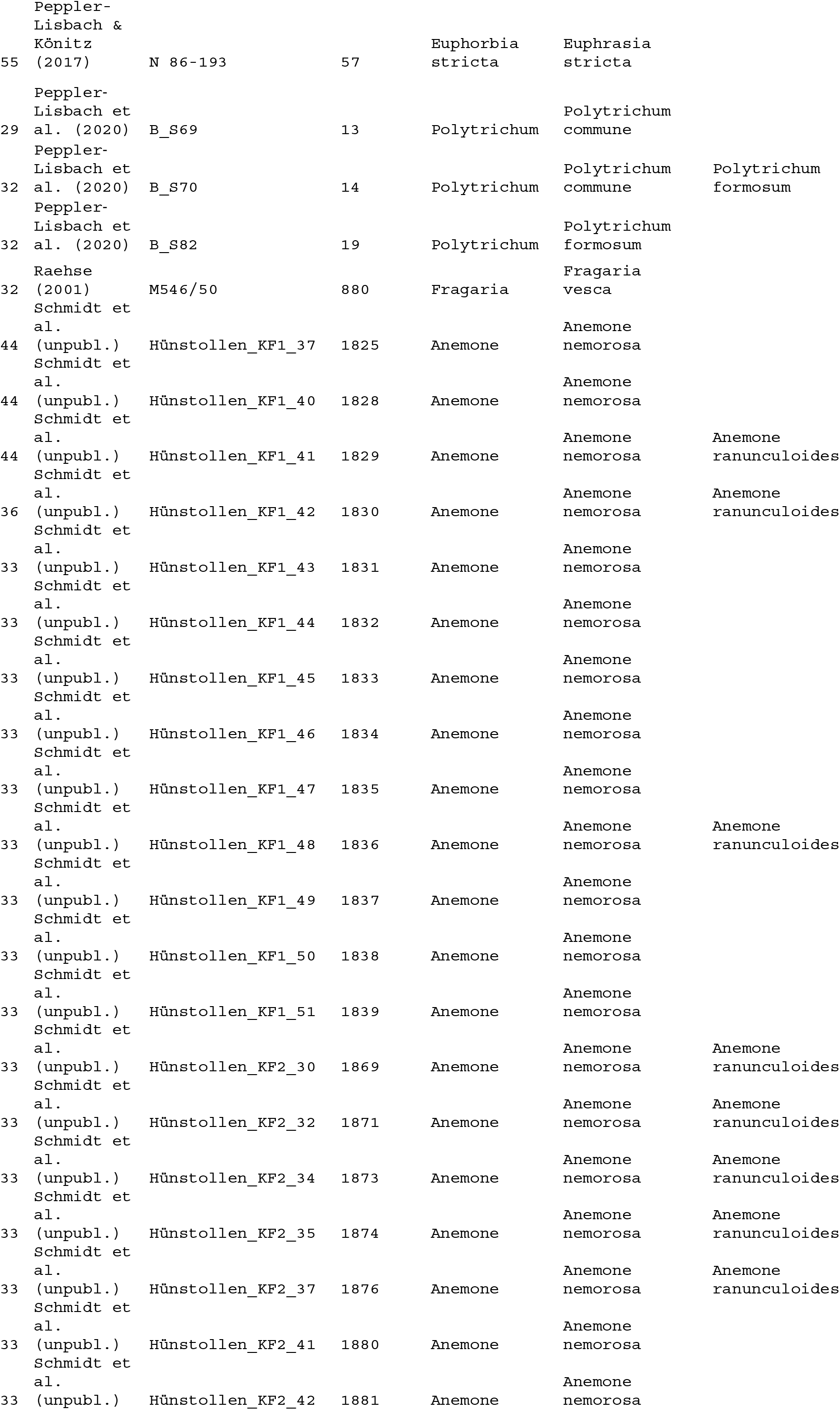
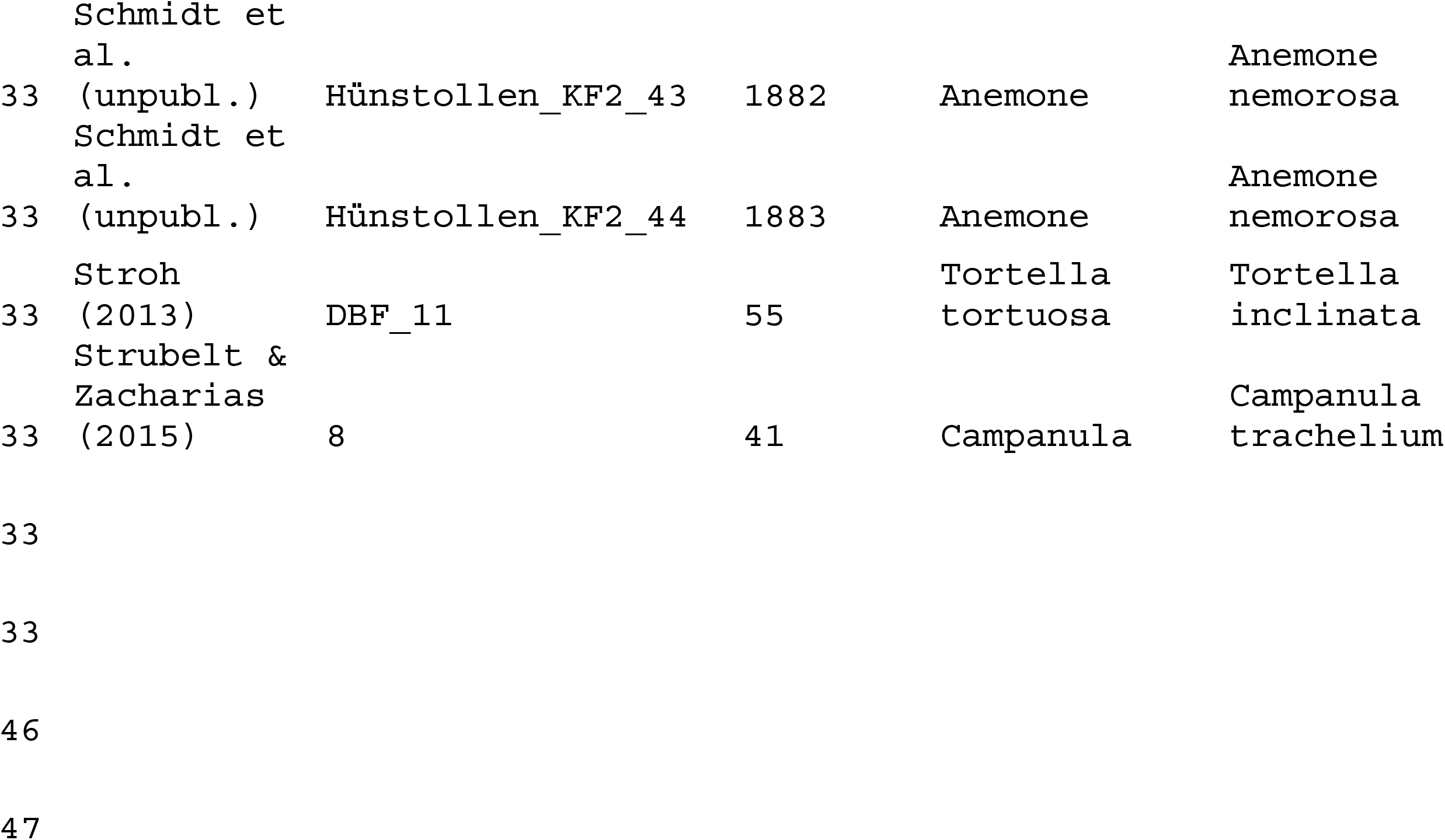
List of all taxon names that were adapted within projects, in addition to the harmonisation across all projects, as shown in Table 4. PROJECT_ID and Project_Name refer to the project in Table 1, RS_PLOT is the plot resurvey ID, which identifies the groups of plots compared in time, RELEVE_NR is the plot observation ID in the Turboveg 2 database (Table 4). Taxon_name_old is the name given by the original author, while Taxon_name_new_1 and Taxon_name_new_2 refer to newly assigned taxon names. In case of two new names the cover values of the old taxon were equally divided among the two new taxa.

The percentage cover values of the same aggregated taxon name of the same plot were merged, assuming a random overlap of their cover values and making sure that the combined cover values cannot exceed 100%^78^. As not all projects had recorded cryptogams, we removed bryophytes and lichens in all projects, using the vegdata package in R^83^. As a result, the original list of 3,280 taxon names that included bryophytes and lichens was reduced to 1,794 taxon names of vascular plants. However, if data on lichens and bryophytes are required, they are available on request from the respective dataset custodians (see Table 1).

The data structure of the header file of ReSurveyGermany follows the Turboveg 2 standard^80^ and in addition holds the fields of ReSurveyEurope (http://euroveg.org/eva-database-re-survey-europe) (Table 4). The fields relevant for the resurvey are RS_PROJECT, which refers to the resurvey project in Table 1. The header field RS_SITE holds the location name of plots and allows for a local geographical scale aggregation of resurvey plots within projects. LOCALITY provides more details on the locality in German

Within each project, the field RS_PLOT holds a plot resurvey ID that connects plot observations from different times made on the same plot. In resurveys, there are also cases, where the previously provided location was not precise enough. In these cases, resurveys often used several plots to match one previous plot, resulting in a one-to-many relationship. If a set of plots at the same site was compared with plot records from another point in time, this field is empty and the unique identifier indicating which plots have to be matched is found in the field RS_SITE. We still keep the original observation ID that a plot received when it was surveyed (RS_OBSERV). We report the exact DATE when a record was made (if available). In addition, the field YEAR lists the year in which the plot was (re)surveyed. If available, we also report the year of the underlying publication (YEAR_PUBL).

Plot area (SURF_AREA) ranges from 0.5 to 2500 m^2^, with 25, 100 and 400 m^2^ being the most frequently used plot sizes (Fig. 4). Plot sizes larger than 100 m^2^ were typical of forest sites (with a very few exceptions).

**Fig. 4:**
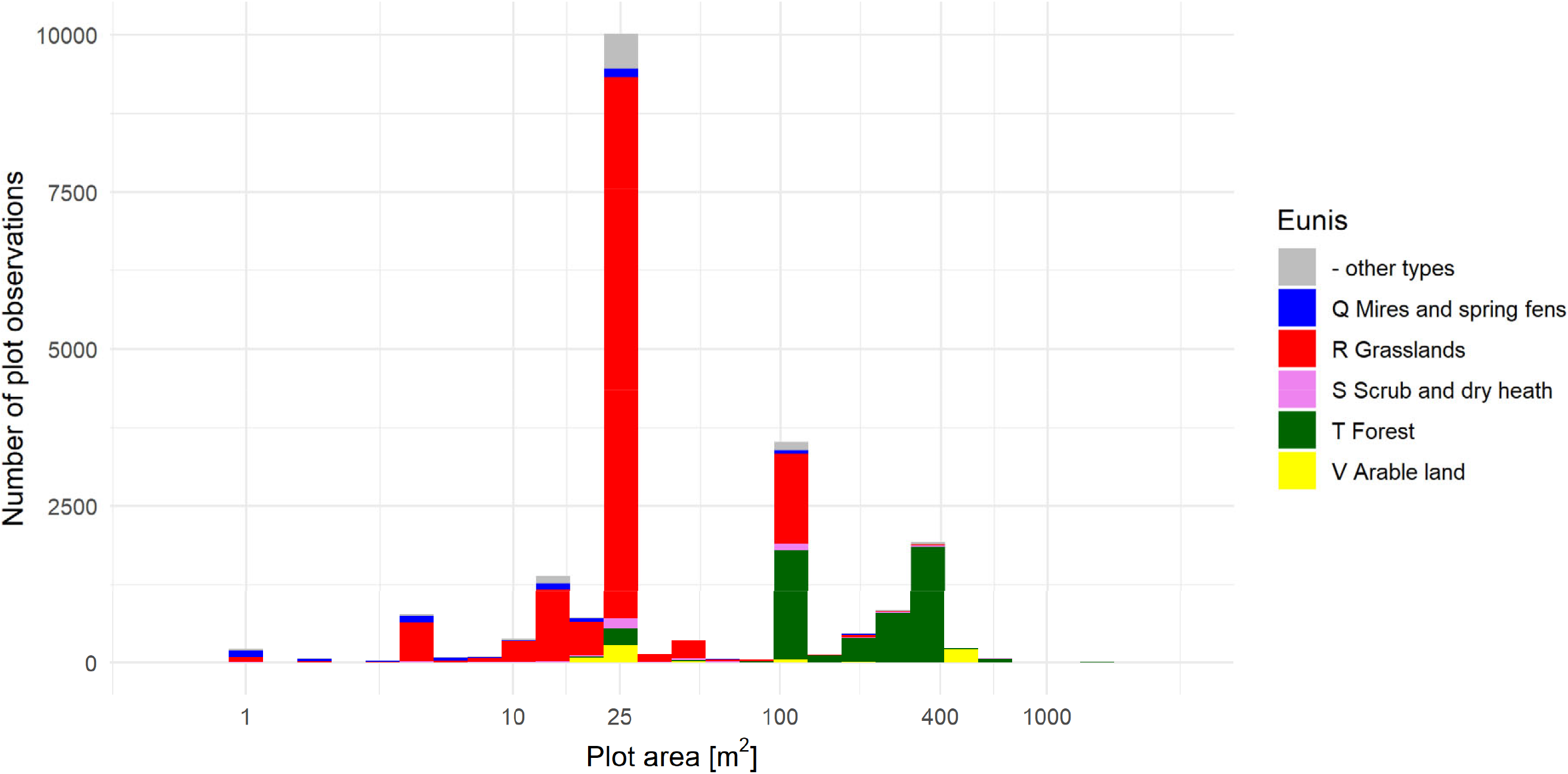
Histogram ofplot size across all records (n=23,641). Colours show Eunis level 1 habitat types.

Geographic information is given by LONGITUDE, LATITUDE and ALTITUDE. Current monitoring programs and data protection of land owners do not allow us to provide location information at the highest available precision. In addition, some records contain occurrence data of rare and protected species. Thus, information on longitude and latitude was rounded to two decimal digits. Compared to the coordinates at highest available precision, rounding resulted in a mean uncertainty of 371 m (± 138 m standard deviation), and thus, is within the somewhat limited range of accuracy provided by many custodians in the first place (see field PRECISION). If more precise coordinates are required for certain analysis we recommend to contact the respective data owners (as shown in Table 1). Vegetation-plot time series differ with respect to the accuracy of the plot relocation during the resurvey. In the ideal case, plots are permanently marked, using poles, metal tent pegs or magnets and metal detectors to retrieve their position (shown as “01” in the LOC_METHOD field, Table 4). In other cases, plots only have exact coordinates (using GPS coordinates, “03” or “04”) or other ways of descriptions of the exact locality (such as from maps, “05”), but are not marked on the ground, which we refer to as semi-permanent plots. In addition, there is information on the cover scale used for the record, a reference to the data source (or, if published, the publication ID), including the table and column from which the data were taken.

The orientation of the plot can be taken from SLOPE (inclination) and slope ASPECT (compass directions). Vegetation structure is described by the height and cover of the different layers, ranging from tree layer to moss layer and including information on cover of litter and bare soil (if available).

Some of our projects included experimental treatments with different management of habitats (e.g. abandonment or establishment of grazing, succession and disturbance). Plots with experimental manipulation contain “Y” in the MANIPULATE) field. The type of manipulation can be taken from MANIPTYPE. When projects involved treatments that are not representative for biodiversity change in the study, we included only the control plots ^44^, plots that reflected the predominant land use at the site (e.g. mowing for a grassland to counteract natural succession) ^20^, that were unfenced ^84^ or were subjected to continuous grazing ^85^.

## Usage notes

The data of the ReSurveyGermany dataset as described above is available https://doi.org/10.25829/idiv.3508-c17blk under the terms specified by CC BY 4.0.

[Please note that the link is not yet activiated, which will happen around May 25^th^, 2022. In the meantime you can already access the metadata via

https://idata.idiv.de/ddm/Data/ShowData/3508?version=0 and the full dataset here: https://cloud.uni-halle.de/s/wei1ljqnq2Wet0A

This part marked in yellow will then be deleted from the paper]

Users are urged to cite the original sources when using ReSurveyGermany in addition to the present paper (see Table 1). As some of the time series will be continued, it might be useful to contact the respective data owners. As described above, the dataset cannot be considered representative of Germany’s vegetation, neither spatially, nor temporally, which is typical of vegetation-plot time series^86^. As plots were established with different objectives in different habitats at different points in time, analysis of vegetation-plot resurveys faces various methodological challenges^60^. Yet, we note that ReSurveyGermany covers about 60% of the 2,988 vascular plant species that occur in Germany (without subspecies and segregates^82^) and includes rare habitats which often harbour rare plant species. This means that even if our sites are not fully representative of the vegetation of Germany and its change over the last century, the data nevertheless give important insights into biodiversity change at the level of local communities and individual species.

## Code availability

The R code to read the plot-species-abundance file (ReSurveyGermany.csv) and combine it with the header data (Header_ReSurveyGermany.csv) is provided on https://github.com/idiv-biodiversity/Read_ReSurveyGermany.

## Acknowledgments

We are grateful to surveyors who recorded vegetation in the field and provided these data. We acknowledge those data contributors who made their data available to us or helped in recording these data: Thea Dittmann, Alexandra Erfmeier, Bernd Gerken, Kerstin Günther, Sabine Heinz, Wilfried Hakes, Heike Heklau, Alfons Henrichfreise, Elisabeth Hüllbusch, Andreas Huwer, Anneke Immoor, Sophie Luise Kühn, Benjamin Krause, Sebastian Leonhardt, Thomas Meineke, Jutta Rach, Jennifer Reinecke, Ulrich Scheidel and Immo Vollmer. We thank Diana Bowler for her analysis of spatial representativeness. The assistance of the iDiv Data & Code Unit (Anahita Kazem, Ludmilla Figueiredo) for archiving the dataset is greatly acknowledged. We very much appreciate the support for the strategic project sMon by the German Centre for Integrative Biodiversity Research (iDiv) Halle-Jena-Leipzig, funded by the German Research Foundation (DFG-FZT 118, 202548816).

